# A comparison of antigen-specific T cell responses induced by six novel tuberculosis vaccine candidates

**DOI:** 10.1101/452060

**Authors:** Miguel J. Rodo, Virginie Rozot, Elisa Nemes, One Dintwe, Mark Hatherill, Francesca Little, Thomas J. Scriba

## Abstract

Eradication of tuberculosis (TB), the world’s leading cause of death due to infectious disease, requires a highly efficacious TB vaccine. Many TB vaccine candidates are in preclinical and clinical development but only a few can be advanced to large-scale efficacy trials due to limited global resources. We aimed to perform a statistically rigorous comparison of the antigen-specific T cell responses induced by six novel TB vaccine candidates and the only licensed TB vaccine, Bacillus Calmette-Guérin (BCG). We propose that the antigen-specific immune response induced by such vaccines provides an objective, data-driven basis for prioritisation of vaccine candidates for efficacy testing. We analyzed frequencies of antigen-specific CD4 and CD8 T cells expressing IFNγ, IL-2, TNF and/or IL-17 from adolescents or adults, with or without *Mycobacterium tuberculosis* (*M.tb*) infection, who received MVA85A, AERAS-402, H1:IC31, H56:IC31, M72/AS01E, ID93+GLA-SE or BCG. Two key response characteristics were analyzed, namely response magnitude and cytokine co-expression profile of the memory T cell response that persisted above the pre-vaccination response to the final study visit in each trial. All vaccines preferentially induced antigen-specific CD4 T cell responses expressing Th1 cytokines; levels of IL-17-expressing cells were low or not detected. In *M.tb*-uninfected and ‐infected individuals, M72/AS01E induced higher memory Th1 cytokine-expressing CD4 T cell responses than other novel vaccine candidates. Cytokine co-expression profiles of memory CD4 T cells induced by different novel vaccine candidates were alike. Our study suggests that the T cell response feature which most differentiated between the TB vaccine candidates was response magnitude, whilst functional profiles suggested a lack of response diversity. Since M72/AS01E induced the highest memory CD4 T cell response it demonstrated the best vaccine take. In the absence of immunological correlates of protection the likelihood of finding a protective vaccine by empirical testing of candidates may be increased by the addition of candidates that induce distinct immune characteristics.

**Author summary:** Tuberculosis (TB) causes more deaths than any other single infectious disease, and a new, improved vaccine is needed to control the epidemic. Many new TB vaccine candidates are in clinical development, but only one or two can be advanced to expensive efficacy trials. In this study, we compared magnitude and functional attributes of memory T cell responses induced in recently conducted clinical trials by six TB vaccine candidates, as well as BCG. The results suggest that these vaccines induced CD4 and CD8 T cell responses with similar functional attributes, but that one vaccine, M72/AS01E, induced the largest responses. This finding may indicate a lack of diversity in T cell responses induced by different TB vaccine candidates. A repertoire of vaccine candidates that induces more diverse immune response characteristics may increase the chances of finding a protective vaccine against TB.

## Introduction

Tuberculosis (TB) is an infectious disease of major global importance. More than ten million people were diagnosed with TB disease in 2016 and 1.67 million died [1]. Current efforts to curb the TB epidemic are insufficient to achieve the 2035 targets set by the World Health Organization, of a 95% reduction in TB deaths and a 90% reduction in the TB incidence rate, compared with levels in 2015 [1]. There is widespread consensus, supported by epidemiological modeling, that a highly efficacious TB vaccine is necessary to achieve these TB control objectives [2, 3].

Thirteen novel TB vaccine candidates were being assessed at various stages in phase 1-3 clinical trials in 2017. However, whilst criteria have been proposed to guide advancement of vaccine candidates to efficacy testing, these are neither unanimously agreed upon nor used [4]. There is also limited stakeholder effort to harmonize or standardize clinical trial design and, as a result, different vaccine candidates are typically assessed in unrelated trials with unique design features that preclude direct comparison of results. This is also the case for assessment of immunological outcomes of TB vaccine trials. Most clinical trials measure vaccine-induced CD4 and/or CD8 cells expressing the Th1 cytokines IFNγ, TNF and IL-2, on the basis that these cells are necessary, although not sufficient, for protective immunity against *Mycobacterium tuberculosis* (*M.tb*) in animal models and humans (reviewed in [5, 6]). However, the types of assays, methodologies and protocols employed by different investigators to measure Th1-cytokine expressing T cells vary considerably [7]. Direct comparison of the magnitude, character and durability of antigen-specific immune responses induced by different TB vaccine candidates is therefore highly problematic. We propose that this is an important gap in knowledge required to guide advancement of vaccine candidates through the clinical development pipeline.

To facilitate rational, data-driven decisions about vaccine candidate advancement, we compared Bacillus Calmette-Guérin (BCG) [8] and six novel TB vaccine candidates, including MVA85A [9, 10], AERAS-402 [11], H1:IC31 [12], M72/AS01E [13, 14], ID93+GLASE [15] and H56:IC31 [16], by their induced antigen-specific CD4 and CD8 T cell responses from data generated in human clinical trials previously completed at the South African TB Vaccine Initiative. The immune responses were measured using the same immunological assay [17, 18], enabling direct comparisons between vaccines. We aimed to define group(s) of vaccines that induced distinct immune responses. Within a group, the vaccines would induce similar immune responses, thus motivating further testing for only one vaccine per group by applying an objective, data-based criterion for vaccine prioritisation.

We identified appropriate statistical approaches and performed an analysis of antigen-specific T cell responses to each antigen in the different vaccine candidates. Comparing the magnitude and cytokine co-expression profiles of vaccine-induced memory CD4 and CD8 T cell responses that persisted to the final study visit in each trial, we revealed considerable homogeneity in vaccine-induced Th1 memory response profiles, particularly in individuals with underlying *M.tb* infection. Our study provides a framework for interpreting immunological characteristics that may be useful for prioritization of vaccine candidates for advancement through the development pipeline.

## Results

Immune responses from a total of 225 (117 *M.tb-*uninfected and 108 *M.tb*-infected) HIV-uninfected individuals vaccinated with one of six novel TB vaccine candidates, and 27 re-vaccinated with BCG, were analyzed (**Tables 1 and 2**). Study participants were either adolescents or adults. Immune responses were measured by a whole blood intra-cellular staining assay (WB-ICS) with multiparameter flow cytometry [17, 18]. For each vaccine, frequencies of CD4 and CD8 T cells that produced IFNγ, TNF, IL-2 and/or IL-17 in response to antigens in that vaccine were measured.

**Table 1.**
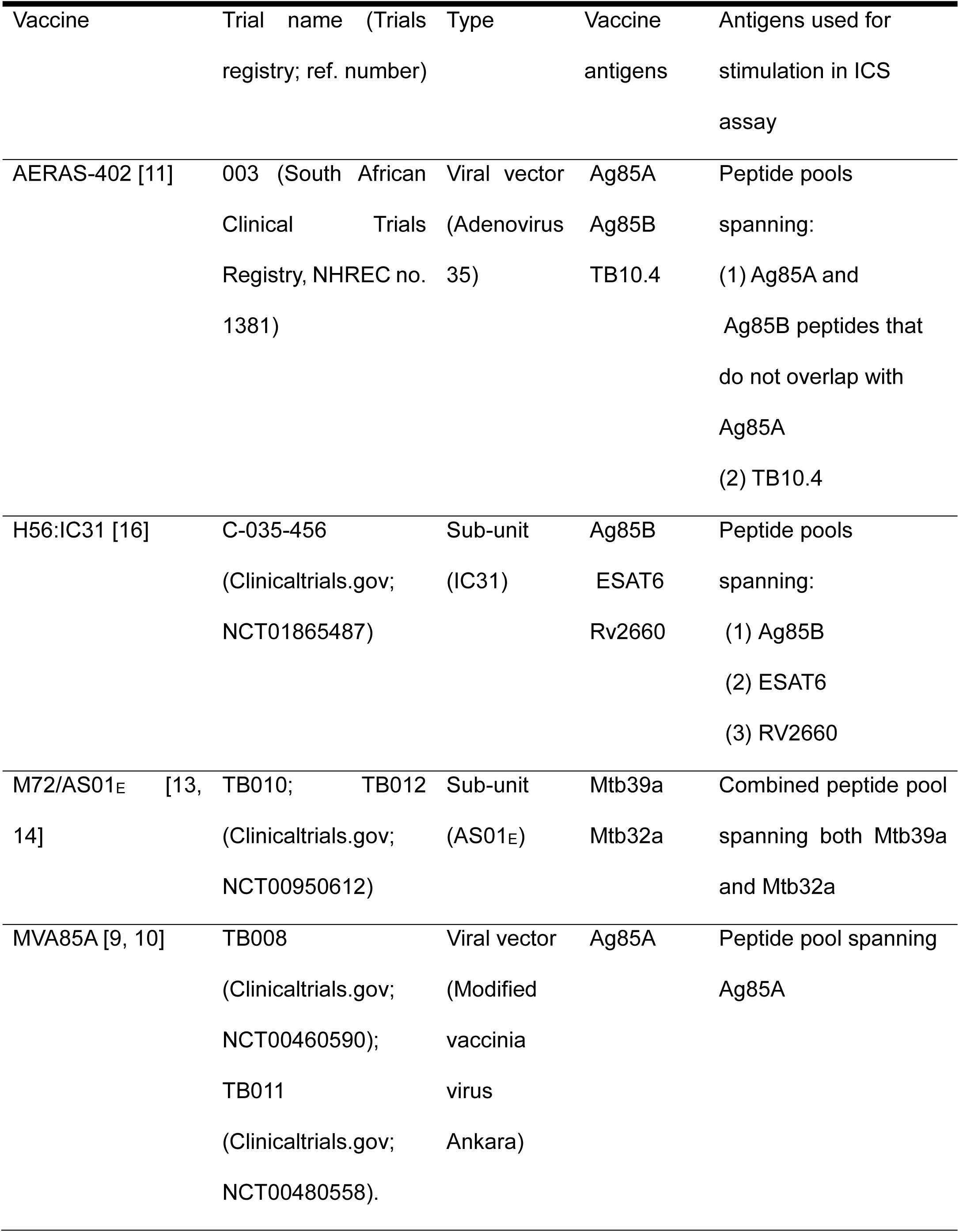
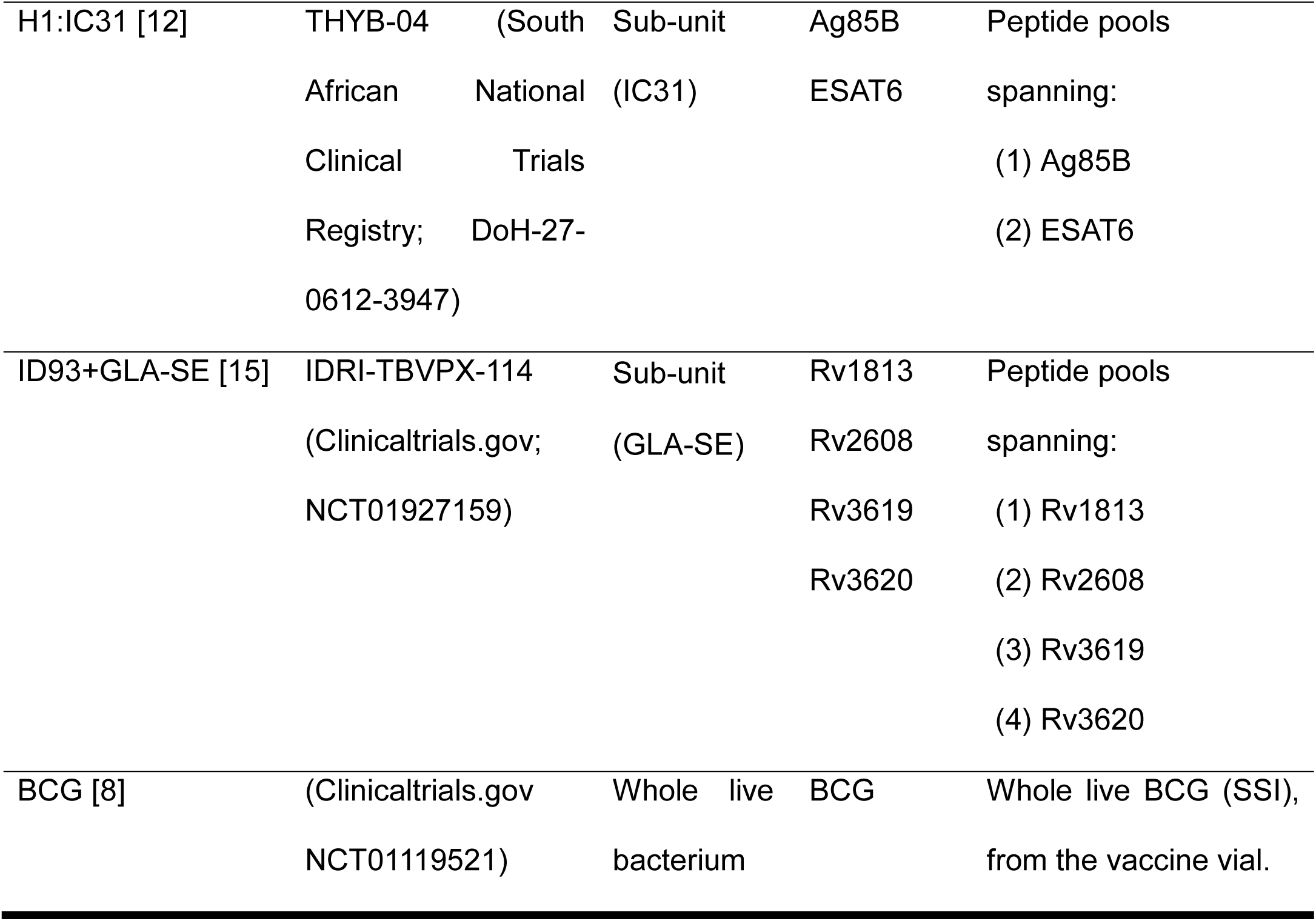
TB vaccine candidates and stimulation antigens.

**Table 2.**
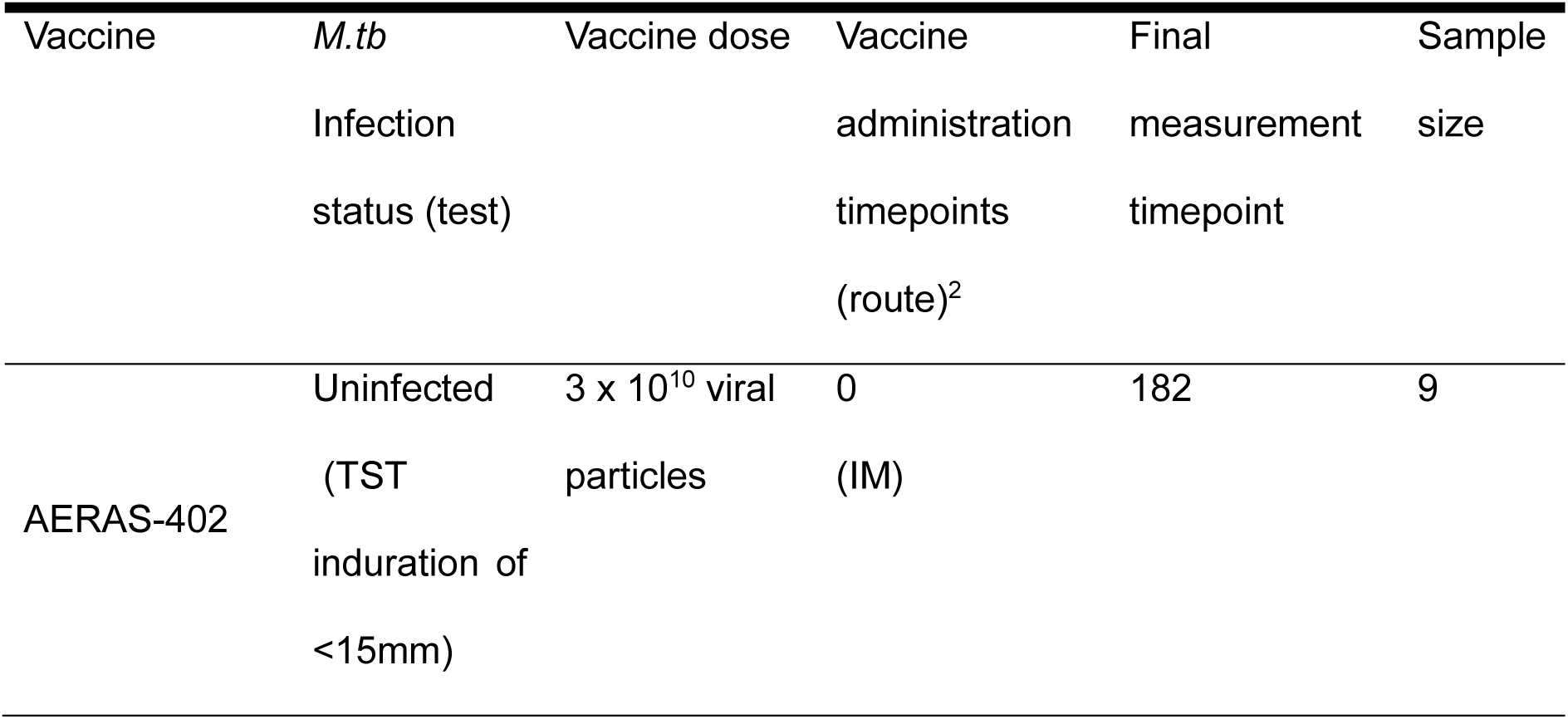
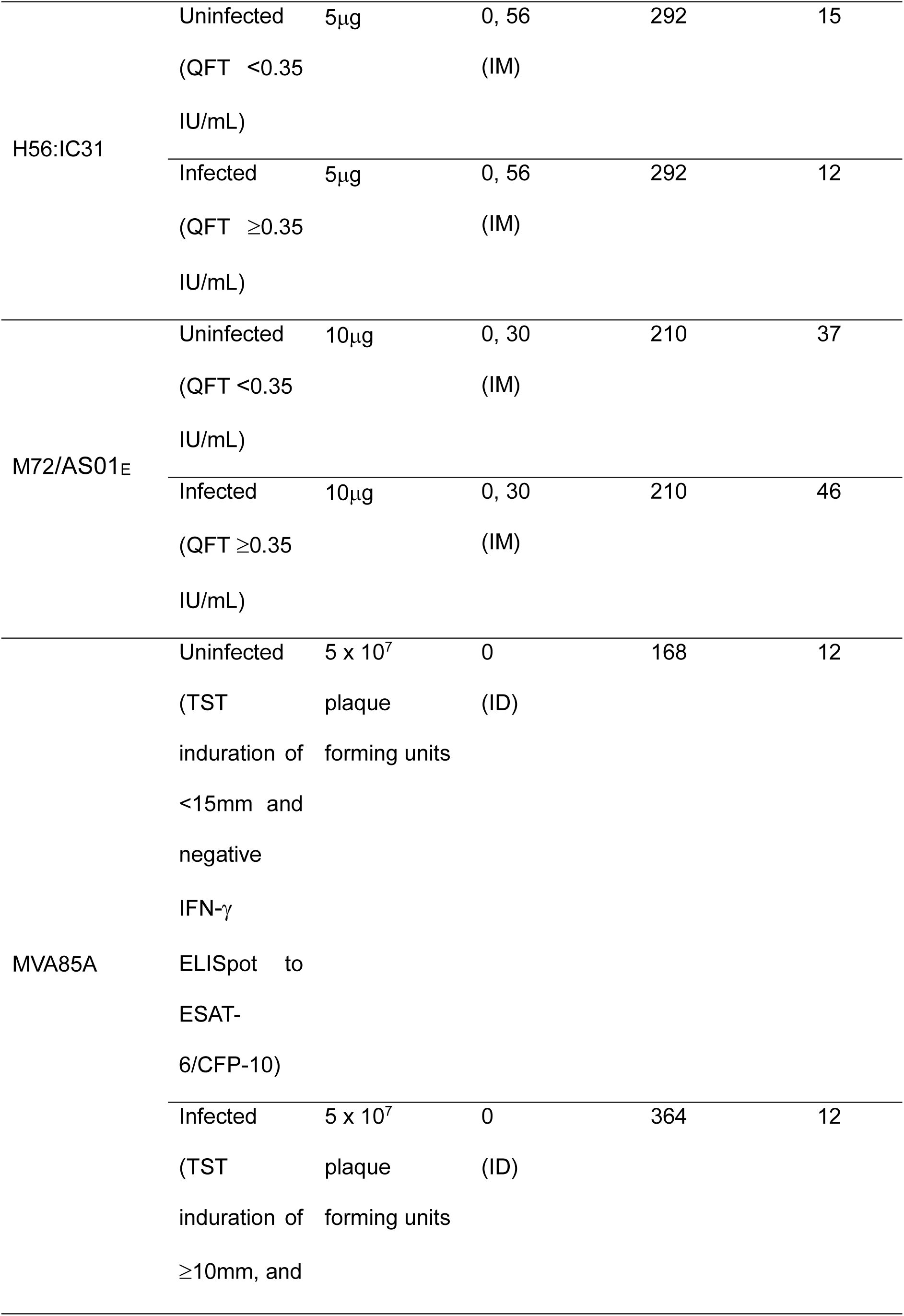
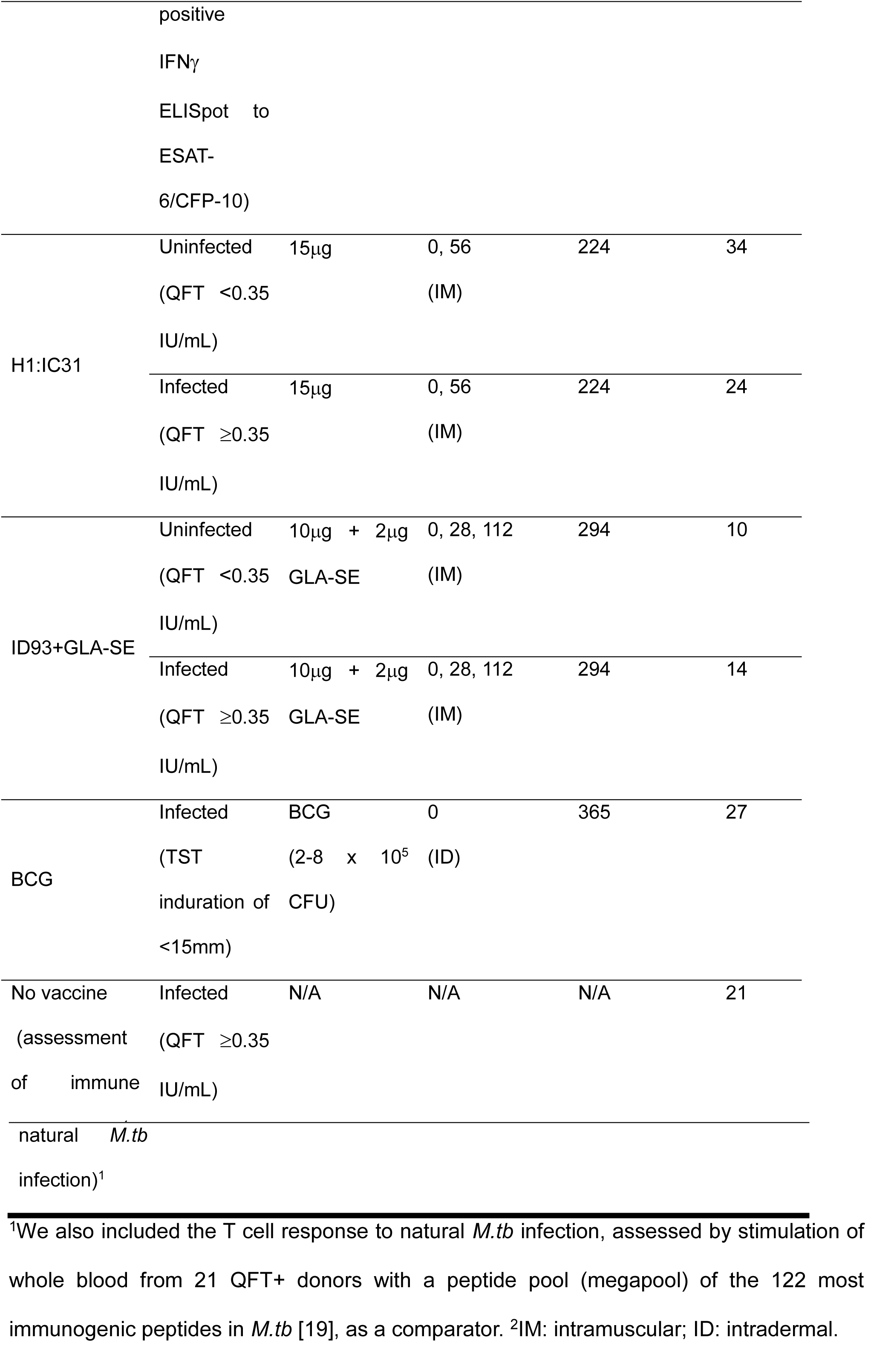
Vaccine groups, administration and measurement schedules and sample sizes.

### Pre-vaccination antigen-specific T cell responses

Many individuals in countries endemic for TB, such as South Africa, are immunologically sensitized to mycobacteria due to a combination of infant BCG vaccination, exposure to environmental non-tuberculous mycobacteria and/or natural *M.tb* infection [20]. To investigate effects of this on pre-vaccination T cell responses, we examined frequencies of Th1-cytokine expressing T cells specific to antigens in each vaccine. CD4 T cell responses to all vaccine antigens were low in *M.tb*-uninfected participants, although responses to antigens in H56:IC31, H1:IC31 and M72/AS01E were detected at frequencies significantly higher than 0.005%, which we defined as the positive response criterion (**Fig 1A** and **S1 Fig**). By contrast, pre-vaccination CD4 responses to antigens in AERAS-402, MVA85A and ID93+GLA-SE were not significantly larger than 0.005%.

**Fig 1.**
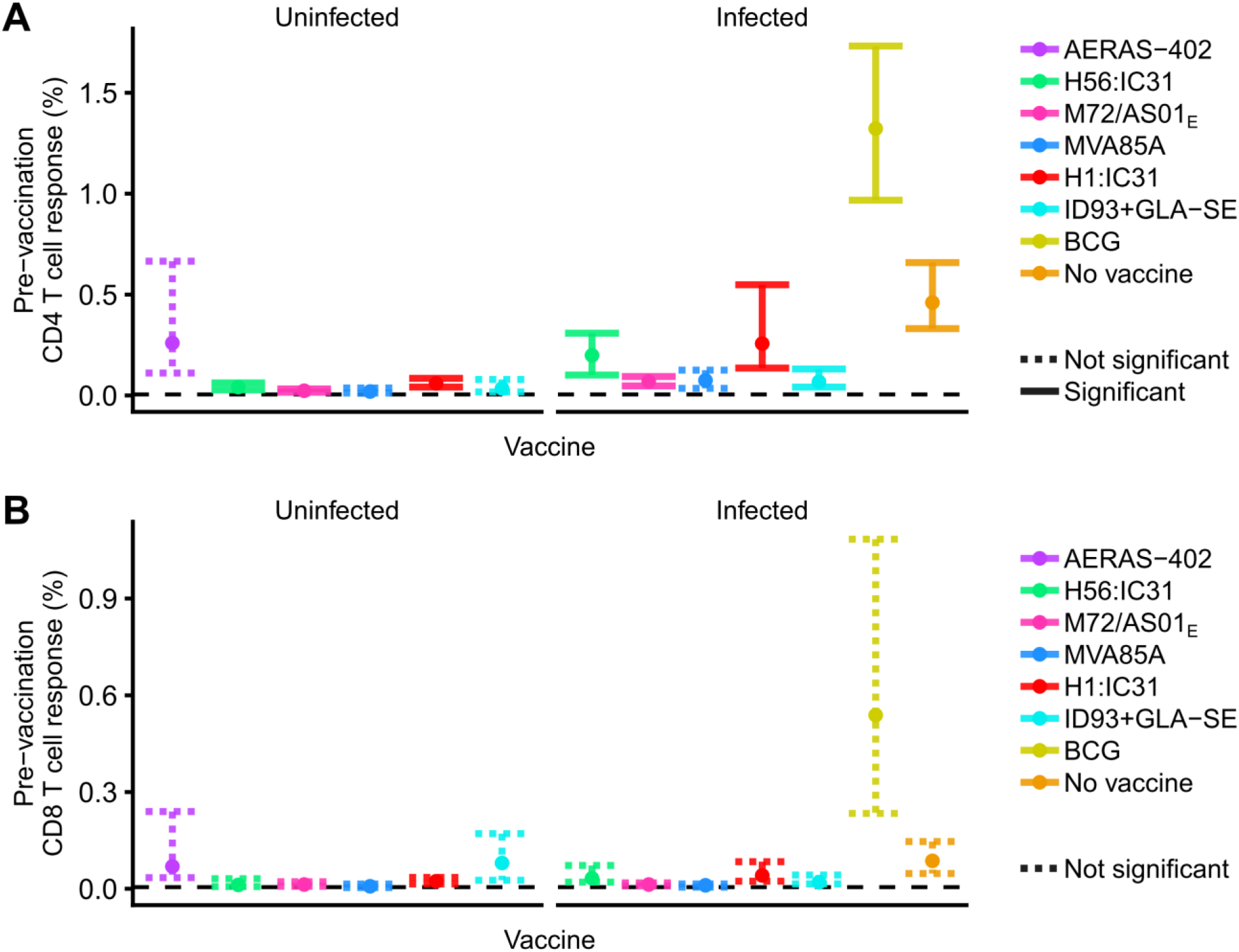
Pre-vaccination antigen-specific CD4 (A) and CD8 (B) T cell responses by vaccine and *M.tb* infection status. Frequencies of antigen-specific, Th1-cytokine expressing CD4 or CD8 T cells pre-vaccination. Points denote sample trimmed means and error bars denote 95% CI. Solid error bar lines indicate responses that significantly exceeded 0.005% after controlling the false discovery rate at 0.01. Dashed lines did not meet this significance criterion. “No vaccine” indicates the immune response to *M.tb* infection detected after megapool stimulation in unvaccinated, IGRA-positive individuals. Pre-vaccination responses to each individual antigen in each vaccine are shown in **S1A Fig**.

In *M.tb*-infected persons, Th1 responses to antigens in all vaccines except MVA85A were detectable at frequencies significantly higher than 0.005%, although the highest responses were observed for BCG and the megapool (**Fig 1A**). The pre-vaccination responses were higher than in *M.tb*-uninfected individuals, but only the difference for M72/AS01E was significant (**S2 Fig**). CD4 T cell responses to Ag85A were not detectable at frequencies significantly exceeding 0.005% (**S1A Fig**).

Pre-vaccination CD8 T cell responses to all vaccine antigens as well as the *M.tb* megapool were low and did not significantly exceed 0.005% for any vaccine in either *M.tb*-uninfected or ‐infected participants (**Fig 1B** and **S1B Fig**).

### Longitudinal vaccine-induced T cell responses

Next, we examined longitudinal frequencies of antigen-specific CD4 and CD8 T cell responses induced by each of the vaccines (**Fig 2**), measured at various time points in each trial (**Table 2**). Vaccine-induced CD4 T cell responses were higher than CD8 T cell responses. In addition, as expected and previously described [8-15, 21], these responses typically peaked at the measurement timepoint immediately after vaccine administration and waned thereafter. Inter-donor variability in CD4 T cell responses to M72/AS01E, H1:IC31 and BCG was high, especially during the early effector phases of the response kinetics (**Fig 2**).

**Fig 2.**
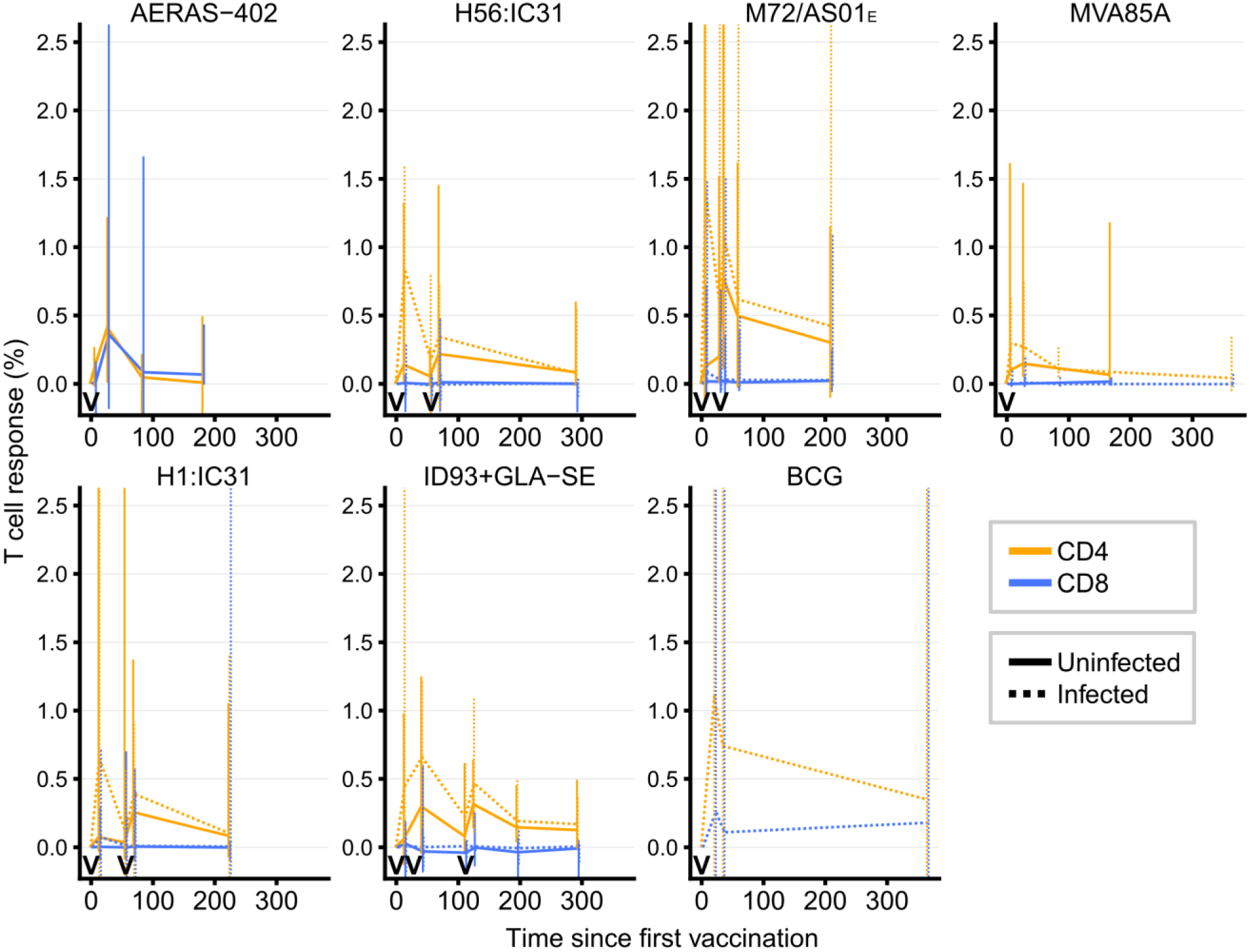
Longitudinal vaccine-induced antigen-specific CD4 and CD8 T cell response kinetics by *M.tb* infection status for the six novel TB vaccine candidates and BCG. Non-vertical lines pass through median CD4 (orange) or CD8 (blue) T cell responses at each timepoint. Vaccine administration timepoints are denoted by the “V” symbols along the xaxes. Vertical solid (*M.tb*-uninfected) and dashed (*M.tb*-infected) lines indicate interquartile ranges. The plots are truncated between −0.1 and 2.5 for readability, suppressing lower and/or upper interquartile ranges for some vaccines for some timepoints.

### Vaccine-induced memory T cell responses

To determine which vaccines induced durable T cell responses and compare the magnitude of these responses, we examined vaccine-induced memory CD4 and CD8 T cell response frequencies. The vaccine-induced memory response is the difference between frequencies of antigen-specific Th1-cytokine producing cells at the final time point and the prevaccination timepoint in each trial.

Among novel vaccine candidates, M72/AS01E, ID93+GLA-SE and H1:IC31 induced CD4 responses that were significantly higher than pre-vaccination levels in both *M.tb*-uninfected and ‐infected individuals, with M72/AS01E inducing greater memory responses than the other novel vaccine candidates (**Fig 3A and S3 Fig**). H56:IC31 induced responses that persisted at levels above those observed pre-vaccination only in *M.tb*-uninfected individuals, and no durable response was detected for either AERAS-402 or MVA85A. BCG induced a highly variable response, which also persisted at levels above those observed prevaccination in *M.tb*-infected individuals. There was negligible statistical support for an effect of underlying *M.tb* infection status on the induced memory CD4 T cell response (**S5 Fig**).

**Fig 3.**
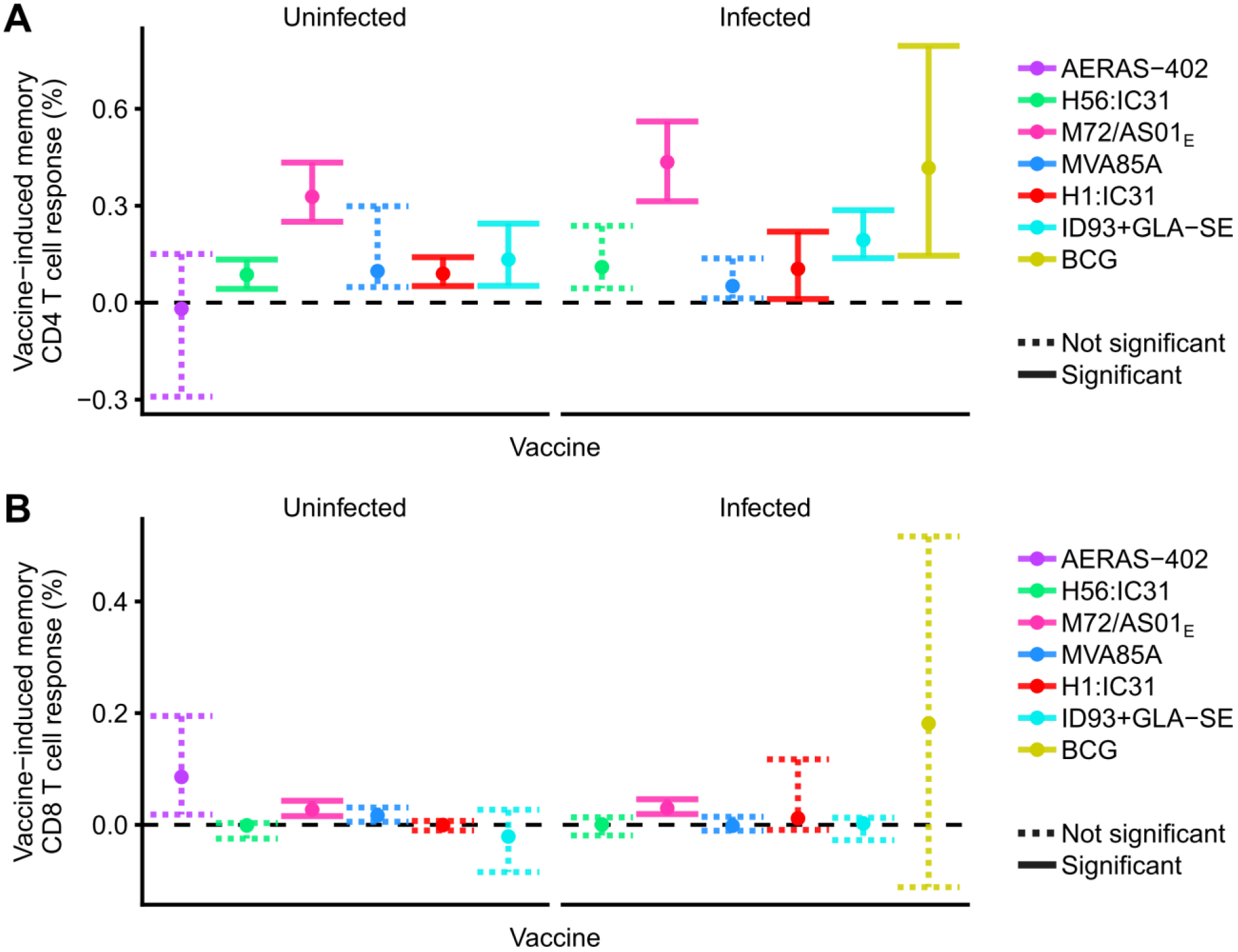
Vaccine-induced memory CD4 (A) and CD8 (B) T cell responses by vaccine and *M.tb* infection status. Frequencies of antigen-specific Th1-cytokine expressing CD4 (a) or CD8 (B) responses at the final time point in each trial, relative to the pre-vaccination frequencies (i.e. memory response minus pre-vaccination response). Points denote sample trimmed means, and error bars 95% CI. Solid error bar lines indicate responses that significantly exceeded 0% after controlling the false discovery rate at 0.01. Dashed lines did not meet this significance criterion.

We also evaluated memory CD4 T cell responses to each individual antigen in each vaccine candidate (**S4A Fig**). Memory CD4 responses induced by H1:IC31 and H56:IC31 in *M.tb*uninfected individuals primarily comprised Ag85B-specific CD4 T cells, whilst a memory response to Rv2608 was predominant in the ID93+GLA-SE-induced response (**S4A Fig**). Vaccine-induced memory CD4 T cell responses to most other individual antigens, except for Mtb32 and Mtb39 (which were not tested separately), were not detected.

M72/AS01E induced low but durable memory CD8 T cell responses in *M.tb*-uninfected and ‐infected individuals. No durable memory CD8 T cell responses were detected for the other vaccines (**Fig 3B**). However, the point estimate of CD8 T cell responses induced by Aeras402 and BCG were higher than M72/AS01E, although this difference was not significant.

Results were similar for memory CD8 T cell responses to each individual antigen in each vaccine candidate; only the combined CD8 response to Mtb32 and Mtb39 significantly exceeded the pre-vaccination response (**S4B Fig**).

Frequencies of vaccine-induced CD4 and CD8 T cells producing IL-17 were low across vaccines, and usually not significantly higher than pre-vaccination levels (**S6 Fig**).

### Cytokine co-expression profiles of vaccine-induced memory responses

An important feature of T cell responses to pathogens, including *M.tb*, is the cytokine co-expression profile of CD4 T cells, which may reflect the degree of T cell differentiation or quality of the response [6, 22, 23]. We analyzed cytokine co-expression profiles of the memory CD4 T cell responses induced by the vaccines. We did not include the cytokine co-expression profiles of the vaccine-induced memory CD8 T cell responses, becasue the magnitudes of the vaccine-induced CD8 memory response were either very low or not significant.

We analyzed cytokine co-expression profiles using principal components analysis (PCA) biplots. In addition, 95% bivariate confidence areas of the mean response for each vaccine were computed using bootstrapping for the first two principal components of the vaccine-induced memory response. In *M.tb-*uninfected individuals, the responses induced by the two viral-vectored vaccines, AERAS-402 and MVA85A, were distinct. This suggests different cytokine co-expression profiles from the four sub-unit vaccines, namely H56:IC31, H1:IC31, M72/AS01E and ID93+GLA-SE. These grouped together regardless of their antigens (**Fig 4A**). IFNγ+ TNF+IL-2+, TNF+IL-2+ and IL-2+ CD4 T cells drove most variation in the first two principal components (**S7A Fig**). The biplot axes (**Fig 4A**) show that MVA85A vaccination was characterized by higher IFNγ+TNF+IL-2+ CD4 T cell responses, the protein sub-unit vaccines by higher TNF+IL-2+ and IL-2+ CD4 T cells, and AERAS-402 by low responses for all cytokine-expressing subsets.

**Fig 4.**
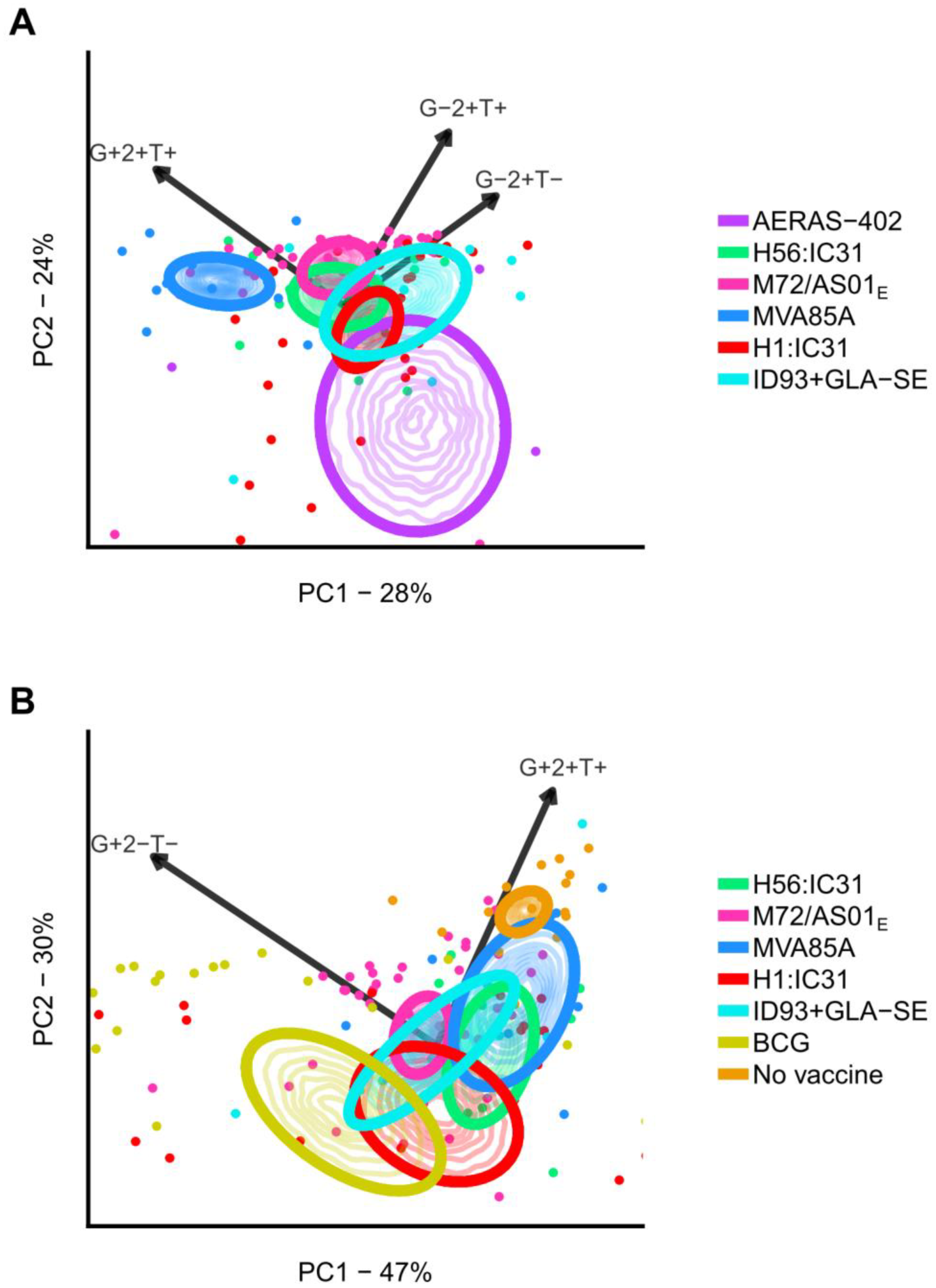
PCA biplots of cytokine co-expression profiles for vaccine-induced memory CD4 T cells. Characterization of vaccine-induced memory CD4 T cells responses by their Th1 cytokine co-expression profiles for each vaccine in *M.tb*-uninfected (A) and ‐infected (B) individuals. PCA biplots show principal components 1 and 2, computed from the scaled vaccine-induced memory T cell responses by cytokine-expressing subset. The scaled response indicates the relative proportions of cytokine co-expressing subsets of the induced response. It ranges from −1 to 1 and is independent of the overall vaccine-induced response magnitude (see Materials and Methods for details). Thick curves denote 95% bootstrap-based confidence areas for bivariate means for each vaccine-induced response. Thin curves represent contour lines of the bootstrap kernel density. The cytokine coexpression combinations displayed (G+2+T+, for example, refers to IFNγ+IL-2+TNF+) by biplot axes had high axis predictivity values relative to other cytokine combinations (**S6 Fig**). Percentages of total variation captured by principal components 1 and 2 are given on plot axes. Points denote observations; not all observations are shown to highlight the confidence areas. The legend item “No vaccine” indicates the group of *M.tb*-infected individuals that did not receive a vaccine, but whose blood was stimulated with megapool.

In *M.tb*-infected individuals, CD4 T cell responses induced by all novel TB vaccines grouped together (**Fig 4B**). Furthermore, the first two principal components captured 77% of the variation, indicating that most variation was due to donor response variability rather than differences in response profiles between vaccines. Variability was largest for IFNγ+TNF+IL-2+ and IFNγ+ CD4 T cell responses (**S7B Fig**). The response to megapool stimulation, induced by *M.tb* infection, was characterized by a predominance of IFNγ+TNF+IL-2+ polyfunctional CD4 T cells, whereas the BCG-induced response comprised low proportions of polyfunctional CD4 T cells; responses induced by the novel TB vaccines fell in between the megapool and BCG responses. Univariate analysis of the responses by cytokine combination for each vaccine corroborated these findings (**Fig 5** and **S8 Fig**).

**Fig 5.**
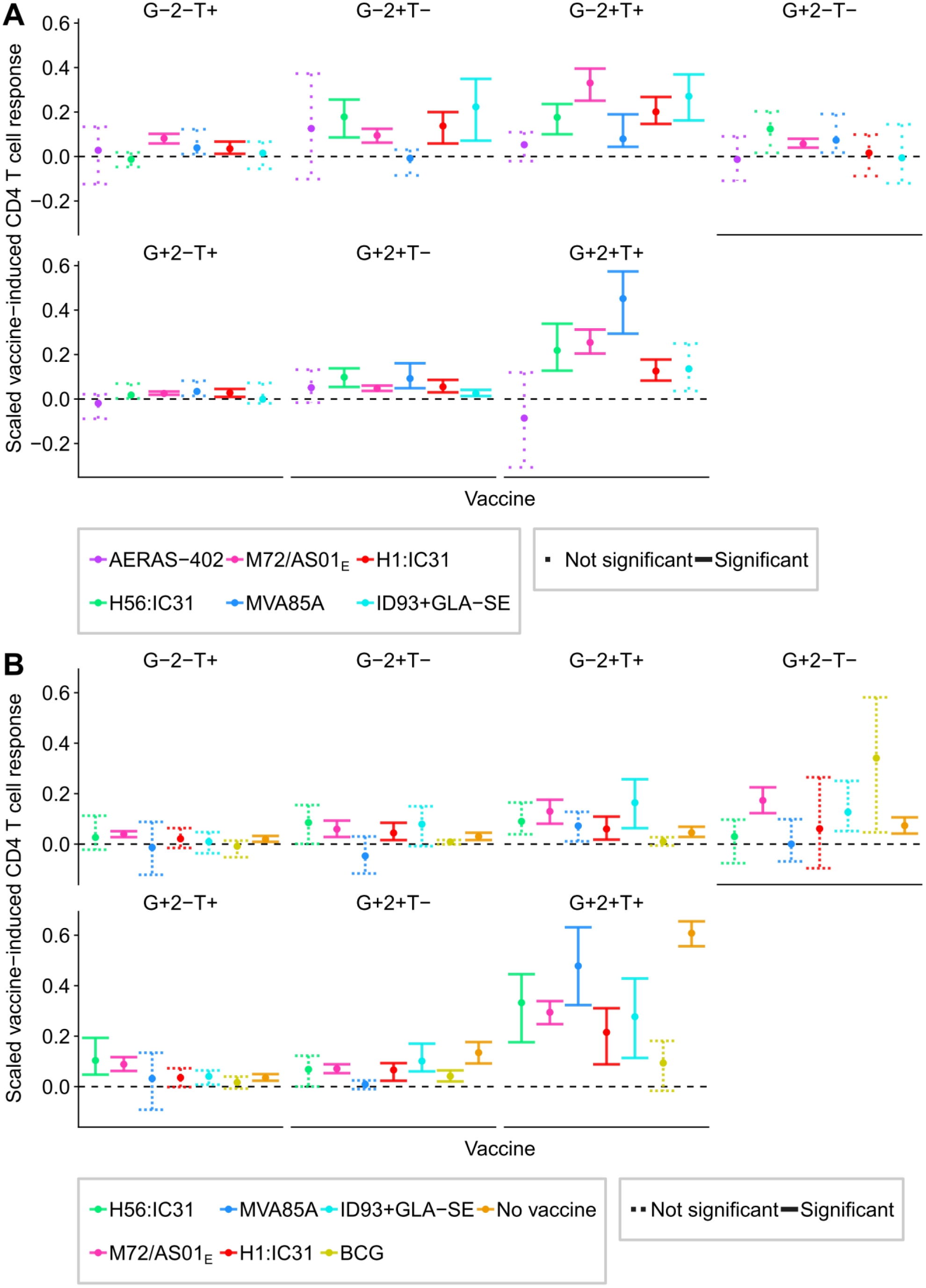
Vaccine-induced cytokine co-expression profiles for vaccine-induced memory CD4 T cells. Characterization of vaccine-induced memory CD4 T cells response by their Th1 cytokine coexpression profiles for each vaccine in *M.tb*-uninfected (A) and ‐infected (B) individuals. Points denote sample trimmed means of the scaled vaccine-induced memory CD4 response for each vaccine for each cytokine-expressing subset, and error bars 95% CI (see Materials and Methods for details). Solid error bar lines indicate responses that significantly exceeded 0% after controlling the false discovery rate at 0.01. Dashed lines did not meet this significance criterion. G+2+T+, for example, refers to IFNγ+IL-2+TNF+.

We also assessed the effect of underlying *M.tb* infection on cytokine co-expression profile (**S9 Fig**). We examined in a univariate analysis differences in scaled vaccine-induced memory responses of TNF+IL-2+, IFNγ+ CD4 T cells and IFNγ+TNF+IL-2+ polyfunctional CD4 T cells between *M.tb*-uninfected and ‐infected persons. The results suggest that the uniformity of response profiles observed in *M.tb*-infected individuals was driven by the protein sub-unit vaccines inducing less TNF+IL-2+ CD4 T cells, and possibly more IFNγ+TNF+IL-2+ CD4 T cells, in *M.tb*-infected individuals.

Taken together, these analyses show that the six novel vaccine candidates induced similar CD4 memory response profiles, exacerbated by a reduction in TNF+IL-2+ CD4 T cells in persons with underlying *M.tb* infection.

## Discussion

TB vaccine development has substantially progressed in recent years, with great advances in our understanding of vaccine platforms, antigen and adjuvant selection and correlates of risk of TB disease [4]. Animal models have been standardized to enable head-to-head comparison and selection of candidate TB vaccines for advancement to phase I clinical trials [4]. Thirteen TB vaccine candidates were assessed in clinical trials in 2017 [4], but only one or two can be advanced to large-scale efficacy trials due to limited global resources.

To provide a data-driven basis for selection of vaccine candidates for further testing in efficacy trials, we performed a comparison of antigen-specific T cell responses induced by six novel TB vaccine candidates that have been assessed in phase 1b or 2a trials at SATVI. Three major points emerged from our study: (1) Antigen-specific T cell responses induced by the candidate TB vaccines were strongly CD4 T cell biased and predominantly expressed Th1-cytokines, (2) Th1 cytokine co-expression profiles of vaccine-induced memory CD4 T cells, a feature of T cell differentiation and functional quality, demonstrated considerable homogeneity between the vaccine candidates, (3) Analysis of T cell response magnitudes showed that amongst the novel vaccine candidates, M72/AS01_E_ induced the highest memory cytokine-expressing CD4 T cell responses.

Our finding that vaccine-induced responses were strongly CD4 T cell-biased with little IL-17 production, and that Th1-cytokine expression profiles were similar across vaccine candidates, highlights a lack of diversity in immunological responses typically analysed in TB vaccine immunogenicity assessments. This result is perhaps not surprising given that most current TB vaccine candidates were designed to specifically target induction of IFNγ-expressing CD4 T cells, predicated on the well-established evidence that Th1 cells are necessary, although not sufficient, for protective immunity against *M.tb*, based on animal models and human studies(reviewed in [5, 6, 24]). We acknowledge that analysis of Th1-cytokine and IL-17 expressing CD4 and CD8 T cell responses may miss important T cell functions.

We did not include analysis of immunogenicity data from trials performed in age groups other than adults or adolescents, or from trials performed in current or prior TB patients. We also excluded data for other vaccine candidates assessed in clinical trials at SATVI, such as MTBVAC, VPM1002 and H4:IC31, because data from vaccinated adults or adolescents at the end of study time point were not available. The whole live mycobacterial vaccines, MTBVAC and VPM1002, are known to induce responses by a broader range of immune cells [25, 26] than those we observed and might add to the diversity of the immune responses induced by vaccine candidates. Further, we did not include analyses of antigen-specific antibody responses, which may also be important in immunity against *M.tb* [27, 28], largely because antibody responses were not measured in each trial assessed here. It should be noted that high-level antigen-specific IgG responses were induced by a number of these TB vaccine candidates [14, 15] and we suggest that such responses should be measured in vaccine trials and included in future head-to-head comparisons.

Our study revealed negligible evidence of an effect of underlying *M.tb* infection on the vaccine-induced memory response magnitude of CD4 or CD8 T cells for any vaccine, but strong evidence for an effect of *M.tb* infection on the vaccine-induced memory response cytokine co-expression profile. For CD4 T cells, *M.tb* infection was associated with a reduction in TNF+IL-2+ and possibly IL-2+ CD4 T cells, which corresponded to an increase in IFNγ+ CD4 T cells for M72/AS01E and possibly IFNγ+TNF+IL-2+ for all novel vaccine candidates. The net effect was that the response profiles induced by MVA85A and the protein sub-unit vaccines in *M.tb*-infected individuals were similar. This drove the response closer to that induced by *M.tb* infection, as detected by megapool stimulation. These data suggest that underlying *M.tb* infection can play a strong role in the character of the vaccine-induced T cell response, as noted in published vaccine trials [12, 14, 21].

Since the vaccine-induced memory Th1 cytokine co-expression profiles were similar, only the response magnitude separated vaccine candidates. M72/AS01E induced the largest antigen-specific CD4 T cell responses, with similar responses between other novel vaccine candidates. Therefore, based on CD4 T cell response magnitude, our study suggests that M72/AS01E demonstrated the best vaccine take, providing support for further clinical testing. Non-immunological differences between vaccines, such as manufacturing cost, potential production capacity and ease of logistical arrangements, are also critical factors in decisions of candidate selection for further testing.

It is important to note that measures of antigen-specific T cell responses, such as the ones we analyzed, do not represent known correlates of protection against *M.tb*. Since immune correlates of protection against *M.tb* remain undefined [4, 6, 27], the induced CD4 and CD8 T cell response producing Th1 cytokines can only be considered a measure of vaccine take. The recent demonstration of protection afforded by BCG re-vaccination against sustained *M.tb*-infection [29] presents an opportunity for elucidating immune correlates of vaccine-induced protection against *M.tb*.

We focused our comparative analyses only on the memory T cell response that persisted to the final study visit of each clinical trial for practical reasons. This is partly because some vaccines were administered once and others twice or three times. We acknowledge that restricting our analyses to the memory response at the final study visit ignores the effector response early after vaccination and other phases of the post-vaccination response, which could have provided more heterogeneity and revealed important immune features for differentiating between the vaccine candidates. However, considering the very different study designs and the fact that the ultimate purpose of vaccination is to induce long-lasting immunological memory, we decided against analysis of earlier time points. Focusing on a single timepoint rather than a longitudinal response also strengthened the statistical analysis. It simplified interpretation and permitted standard multivariate analysis. It also increased statistical power, as it reduced the number of hypothesis tests to perform and assessed the post-vaccination timepoint with the lowest response variability. Moreover, the number of days between the final measurement and both the first vaccination and the last vaccination varied between vaccines, which confounded measurement timings with vaccine. Regardless, it is unlikely that this would have affected our interpretation that M72/AS01_E_ induced the highest T cell response magnitude among novel vaccine candidates, since follow-up time in the M72/AS01E trial was well within the follow-up ranges for the other vaccines. Finally, the small sample sizes for some groups limited our ability to detect significant differences between vaccines.

Our study provides a framework for data-driven vaccine prioritisation. To facilitate these analyses in future, two factors are important. The first is standardization across trials of vaccine candidates, particularly in terms of immunological assay and time points at which immune responses are measured. A common stimulation antigen preparation, such as a “megapool” of *M.tb* peptides (as in [19]), would also facilitate comparisons. The second factor is sample size. Comparing vaccines entails numerous pair-wise comparisons, and increased sample sizes would facilitate detecting and characterising differences.

In conclusion, our study suggests that the T cell response feature which most differentiated between the TB vaccine candidates was response magnitude, whilst functional profiles suggested a lack of response diversity. Since M72/AS01E induced the highest memory CD4 T cell response it demonstrated the best vaccine take. In the absence of immunological correlates of protection the likelihood of finding a protective vaccine by empirical testing of candidates may be increased by the addition of candidates that induce distinct immune characteristics.

## Materials and Methods Novel TB vaccine candidates

The dataset analyzed in this study collates vaccine-specific immune responses from different clinical trials performed at the SATVI Field Site outside Cape Town, South Africa. A summary of the different trials is presented in **Table 1**.

Immune responses were measured by whole blood intra-cellular staining assay (WB-ICS) with multiparameter flow cytometry [17, 18]. Fresh whole blood was stimulated for 12 hours with peptide pools spanning the relevant antigens (**Table 2**), or whole, live BCG.

The study protocol was approved in writing by the Human Research Ethics Committee of the University of Cape Town (HREC ref: 039/2017) and is based on anonymized data from previously published clinical studies.

## Analysis design and assumptions

In some trials different vaccine doses and/or number of administrations were assessed. In these cases, to simplify interpretation of results and increase statistical power, we selected the dose and/or administration strategy that was reported as optimal in the original trial report, based on vaccine safety and tolerability as well as immunogenicity outcomes (**Table 2**). This was generally the dose that induced the highest T cell response magnitude.

As a result of poor standardization between different clinical trials a number of important trial design features differ substantially between the different trials (**Table 2**), including age of vaccinees (ranging from adolescents to adults), method and cut-off for diagnosing *M.tb* infection (tuberculin skin test [TST], ESAT-6/CFP-10 responses detected by IFNγ ELISpot assay or Quantiferon Gold In-Tube [QFT]), number of and timing of vaccine administrations, sampling timepoints for immunological measurements and the duration of participant follow-up.

All novel TB vaccine candidates, except AERAS-402, were given to both *M.tb*-uninfected and ‐infected individuals (**Table 2**). Participants in the AERAS-402 trial were assessed for *M.tb*-infection based on a TST induration of ≥15mm [11]. In the adult MVA85A trials infection was based on a TST induration and a positive response to ESAT-6/CFP-10 peptide pool in an in-house IFNγ ELISpot assay [10] while infection was based on a TST induration of ≥15mm and a positive ELISpot response to ESAT-6/CFP-10 peptide pool in the adolescent MVA85A trial [9]. All other trials used QFT with the manufacturer’s 0.35IU/mL threshold. Comparisons between vaccines were confounded by differences in vaccine-administration schedule (**Table 2**). This study therefore compared overall vaccination strategies, rather than vaccines. Age of participants – adolescents or adults – also varied by vaccine (**Table 2**). Within this study, adolescents and adults were considered immunologically equivalent.

Pre-vaccination is the time point at which the first vaccine was given to an individual. The memory time point for CD4 and CD8 T cell response cytokine expression profiles provides the number of samples after excluding individuals based on negligible change from the prevaccination timepoint. The cut-off used for exclusion was a sum of absolute changes across the different cytokine combinations from pre-vaccination levels of 0.02.

## Immunological measurements

Antigen-specific CD4 and CD8 T cells producing IFNγ, IL-2, TNF and/or IL-17 were measured by WB-ICS assay using flow cytometry as previously described [17, 18]. Frequencies of T cells expressing cytokines in the unstimulated control (background) were subtracted from those in antigen-stimulated conditions; where the background response was greater than the stimulated response, the background-subtracted response was set to zero. When T cell responses for an individual vaccine were measured by separate peptide pools (representing different antigens), background-subtracted response frequencies for the antigens were summed. We defined pre-vaccination responses as the response at day zero (measured before the first vaccine administration) and the memory response as the response at the final time point. To yield the vaccine-induced memory response, we subtracted the pre-vaccination response from the memory response. Where an individual lacked a pre-vaccination response measurement, the median response for its vaccine and *M.tb* infection status group was used.

## T cell response magnitude and cytokine co-expression profile

We analyzed the antigen-specific T cell response magnitude and the cytokine co-expression profile of antigen-specific T cells. To define the response magnitude, let *m_ij_* be the frequencies of vaccine-induced memory CD4 or CD8 T cells for the *j*-th cytokine combination for the *i*-th individual. Then the vaccine-induced response size for the *i*-th individual is equal to

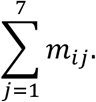

This is the net change from pre-vaccination in frequencies of antigen-specific CD4 or CD8 T cells producing IFNγ, IL-2 and/or TNF.

The cytokine co-expression profile aimed to reflect the T cell differentiation or quality of the antigen-specific T cell response. By using *m_ij_* as defined above, the profile measure for the *i*-th individual for the *j*-th cytokine combination is given by

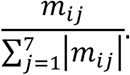

We named this the scaled response. If all the changes from pre-vaccination levels for a certain individual are positive, then the scaled response for a certain cytokine combination gives the proportion of that change that consists of CD4 T cells producing that cytokine combination. The scaled response allows us to compare the extent to which different vaccines “favour” induction of different cytokine combinations, independently of the overall magnitude of the response induced by the vaccine. For analyses of cytokine co-expression profile, we excluded individuals for which the sum of absolute changes from pre-vaccination was less than 0.02, to ensure that the response profile was only analyzed in individuals with memory responses that meaningfully changed relative to pre-vaccination. Number of participants included in this study after exclusion are shown in S1 Table.

## Univariate confidence intervals and hypothesis tests

Because sample sizes were often small (**Table 2**) and the response distributions severely skewed, bootstrapping was used to both construct confidence intervals and perform hypothesis tests for univariate population statistics. Confidence intervals were constructed using the bias-corrected and accelerated method [30] based on 10^4^ bootstrap samples. Hypothesis testing was performed using the bootstrap-*t* approach [31] based on 10^4^ bootstrap samples, with standard errors calculated using the double bootstrap [32] based on 500 bootstrap samples.

We used trimmed means with symmetric trimming of the smallest 20% and the largest 20% of observations, because this is more robust to outliers than non-trimmed means and reflected the typical T cell response better [33]. For one-sample hypothesis tests, the false discovery rate was controlled at 0.01 using the Benjamini-Hochberg procedure [34]. For groups of pair-wise comparisons, the false discovery rate was controlled at an increased 0.05, due to the positive dependency of the tests.

## Multivariate analyses

Multivariate cytokine co-expression profile data were analyzed using principal components analysis (PCA) [35, 36] and biplot axes [37] were calibrated to stretch between the mean and the maximum observed value [38].

To assist interpretation of the biplot, confidence areas for each group were estimated by taking bootstrap samples of the mean, assuming normality of the bootstrap distribution, and applying the fact that the Mahalanoubis distance from the mean has a 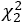 distribution. The normality of the bootstrap samples may be checked by assessing the ellipticity of the contour lines of the bootstrap sample kernel density.

Axis predictivity was used to detect which variables are well represented in a biplot [39] and only variables with high axis predictivity relative to other variables have their axes displayed.

This simplifies biplot interpretation, and indicates which variables primarily drove the variability in the response.

## Acknowledgments

We thank co-investigators and developers/manufacturers of TB vaccine candidates who enabled generation of the data during the original clinical trials. These included Helen McShane and Adrian Hill (University of Oxford), Peter Andersen and Morten Ruhwald (Statens Serum Institute), Paul Gillard and Opokua Ofori-Anyinam (GSK), Steve Reed and Rhea Coler (Infectious Disease Research Institute), Jerald Sadoff, Ann Ginsberg and Tom Evans (Aeras), and Willem Hanekom (SATVI). These individuals and organizations had no role in the analyses presented in this manuscript. Finally, we thank all study participants and the SATVI clinical and laboratory teams.

---

**S1 Fig.**
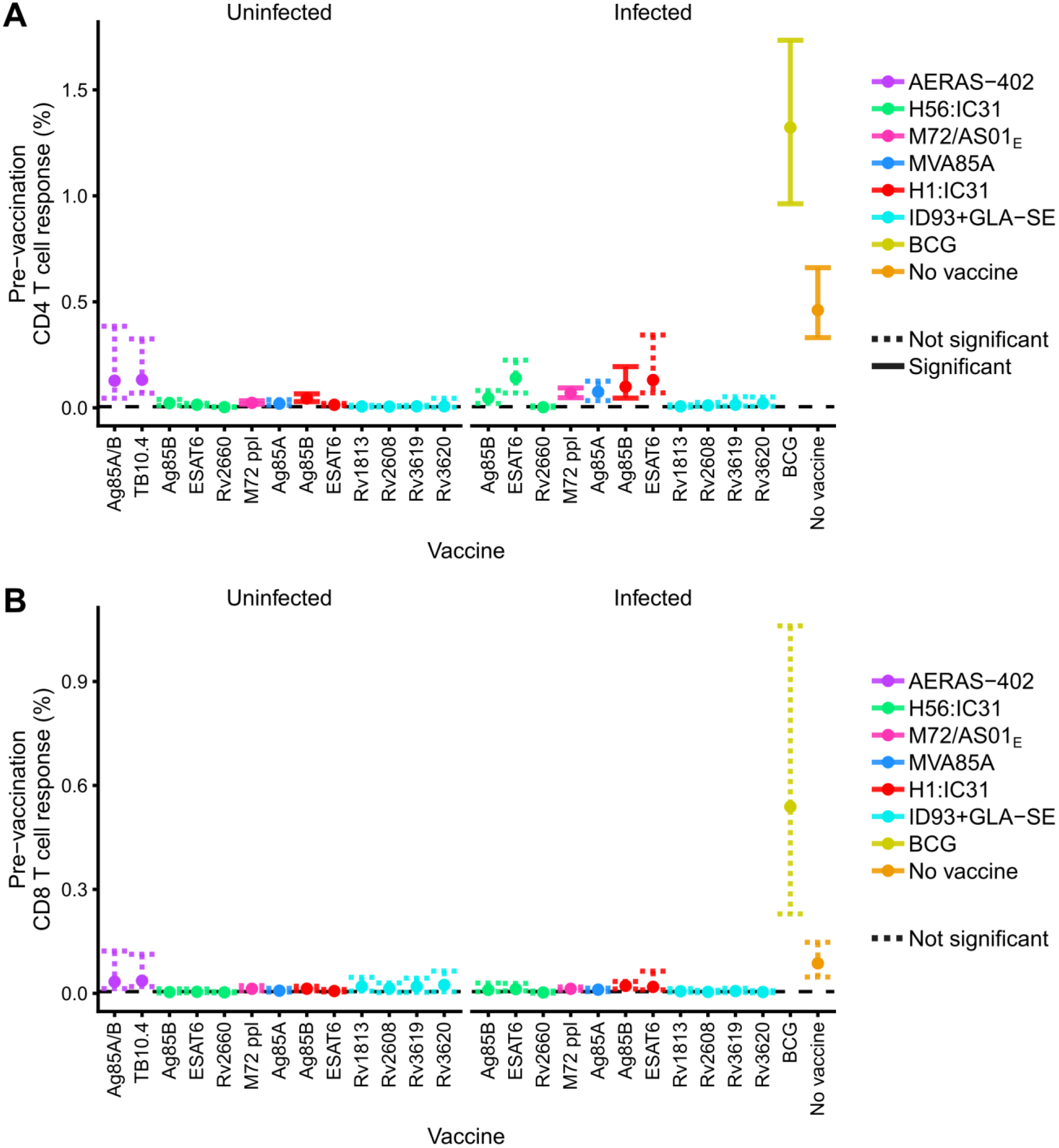
Pre-vaccination antigen-specific CD4 (A) and CD8 (B) T cell responses by individual antigens contained in each vaccine. Frequencies of antigen-specific, Th1-cytokine expressing CD4 or CD8 T cells pre-vaccination. Points denote sample trimmed means and error bars denote 95% CI. Solid error bar lines indicate responses that significantly exceeded 0.005% after controlling the false discovery rate at 0.01. Dashed lines did not meet this significance criterion. “No vaccine” indicates the immune response to *M.tb* infection detected after megapool stimulation in unvaccinated, IGRA-positive individuals.

**S2 Fig.**
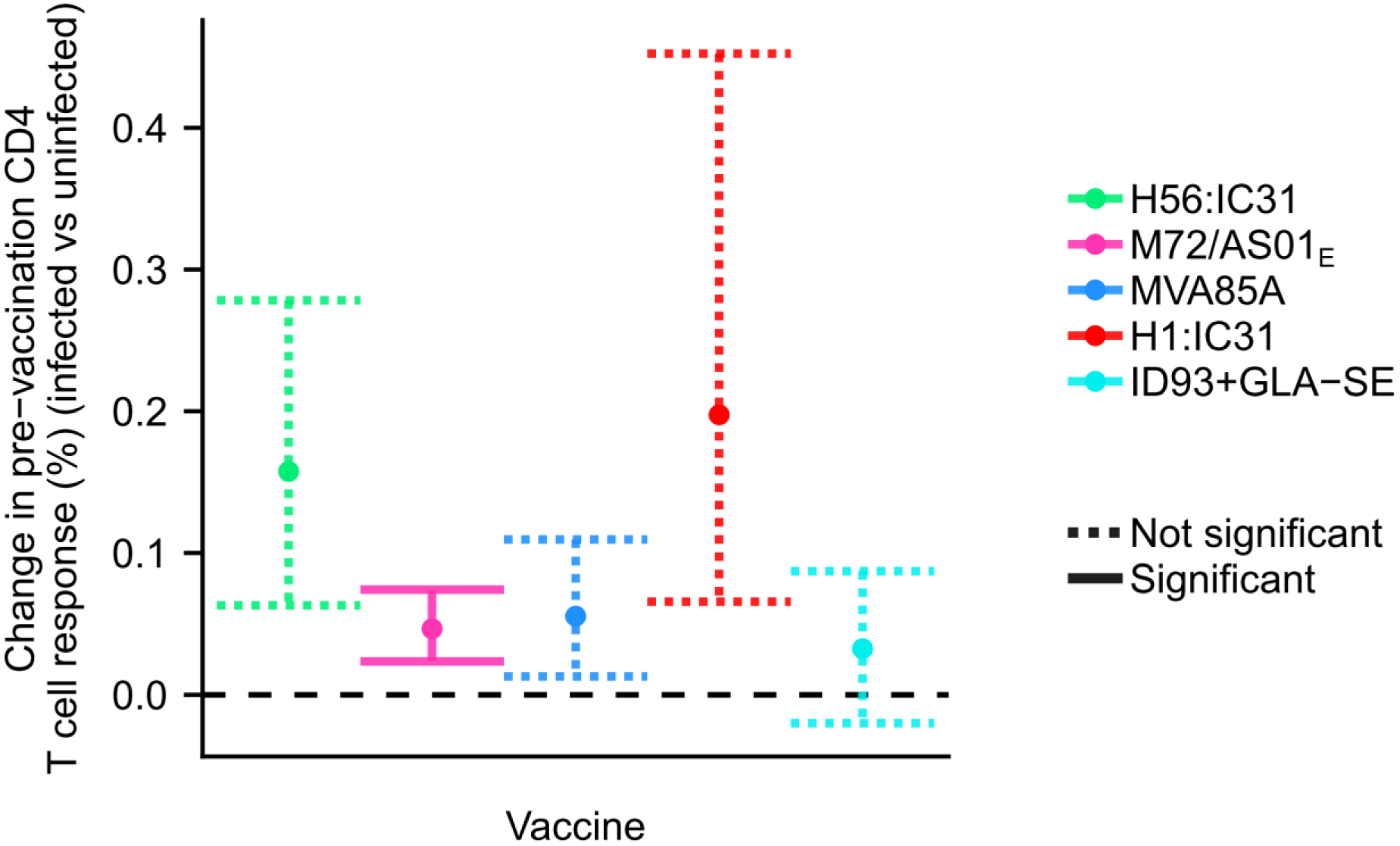
Effect of underlying *M.tb* infection status on pre-vaccination CD4 T cell response magnitudes. Differences between *M.tb*-infected and ‐uninfected individuals in pre-vaccination frequencies of antigen-specific CD4 T cell responses to antigens in each vaccine candidate. Points denote sample trimmed means and error bars denote 95% CI. Solid error bar lines indicate responses that were significantly different between *M.tb*infected and ‐uninfected individuals given the same vaccine, after controlling the false discovery rate at 0.01. Dashed lines did not meet this significance criterion.

**S3 Fig.**
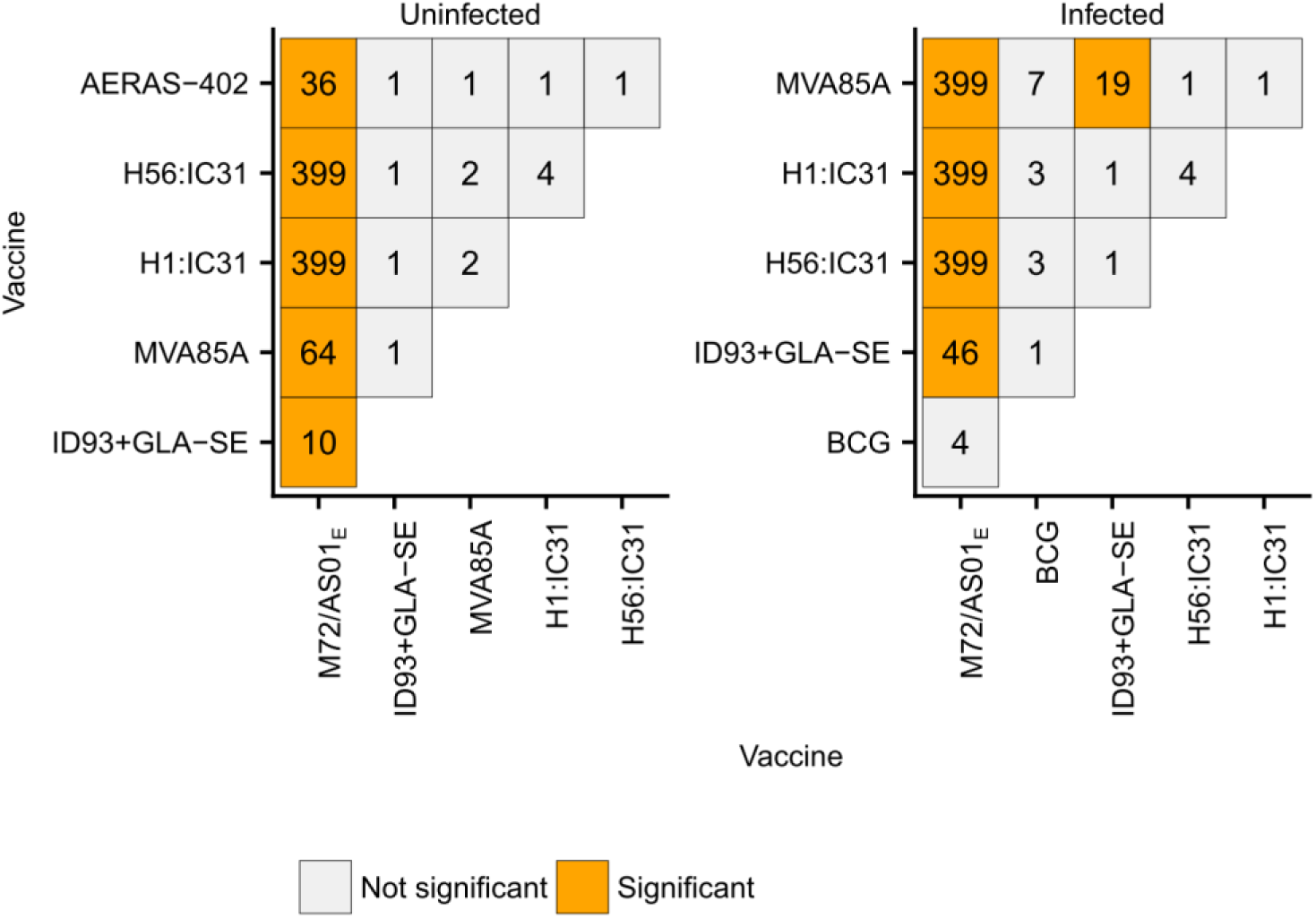
Pair-wise comparisons of vaccine-induced memory CD4 T cell responses between vaccines. Cells display maximum Bayes factors calculated based on the p-values from hypothesis tests for a difference between vaccines in vaccine-induced memory CD4 T cell responses (antigen-specific Th1 cytokine positive CD4 T cells at final trial time point minus pre-vaccination time point). The colour of the blocks indicates statistical significance of the two-sided hypothesis test for a difference in population trimmed means, after controlling the false discovery rate at 0.05. An orange block means that the induced response for the vaccine below the block was significantly larger than the vaccine left of the block. A grey block indicates non-significance. For example, the plot shows that in *M.tb*uninfected individuals the only statistically significant differences were that M72/AS01_E_ induced larger memory responses than all other vaccines.

**S4 Fig.**
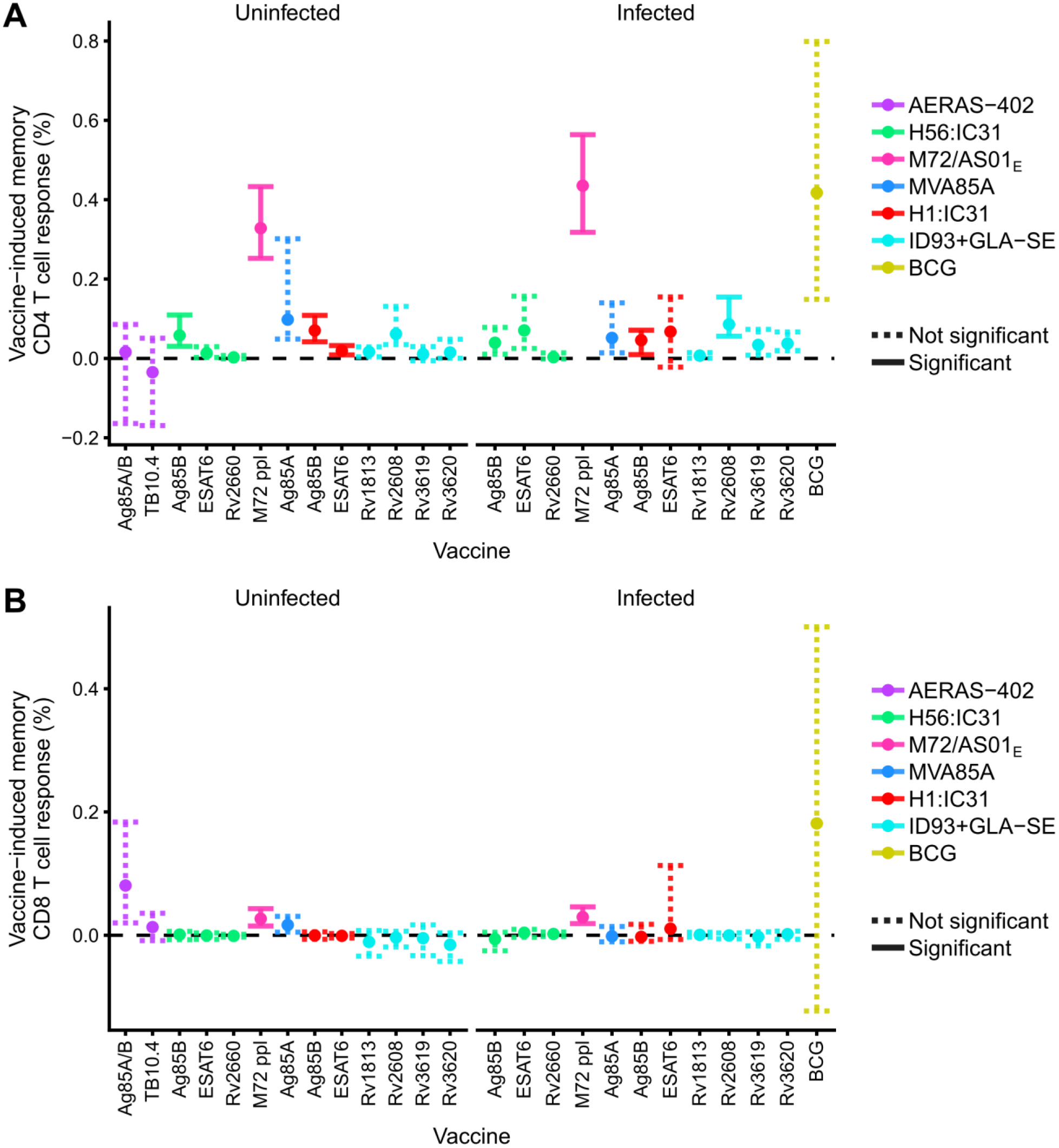
Antigen-specific memory CD4 (A) and CD8 (B) T cell responses by individual antigens contained in each vaccine. Vaccine-induced frequencies of antigen-specific memory CD4 T cell responses to antigens in each vaccine candidate (antigen-specific Th1 cytokine positive CD4 T cells at final trial time point minus pre-vaccination time point). Points denote sample trimmed means and error bars denote 95% CI. Solid error bar lines indicate responses that significantly exceeded 0.005% after controlling for a false discovery rate at 0.01. Dashed lines did not meet this significance criterion.

**S5 Fig.**
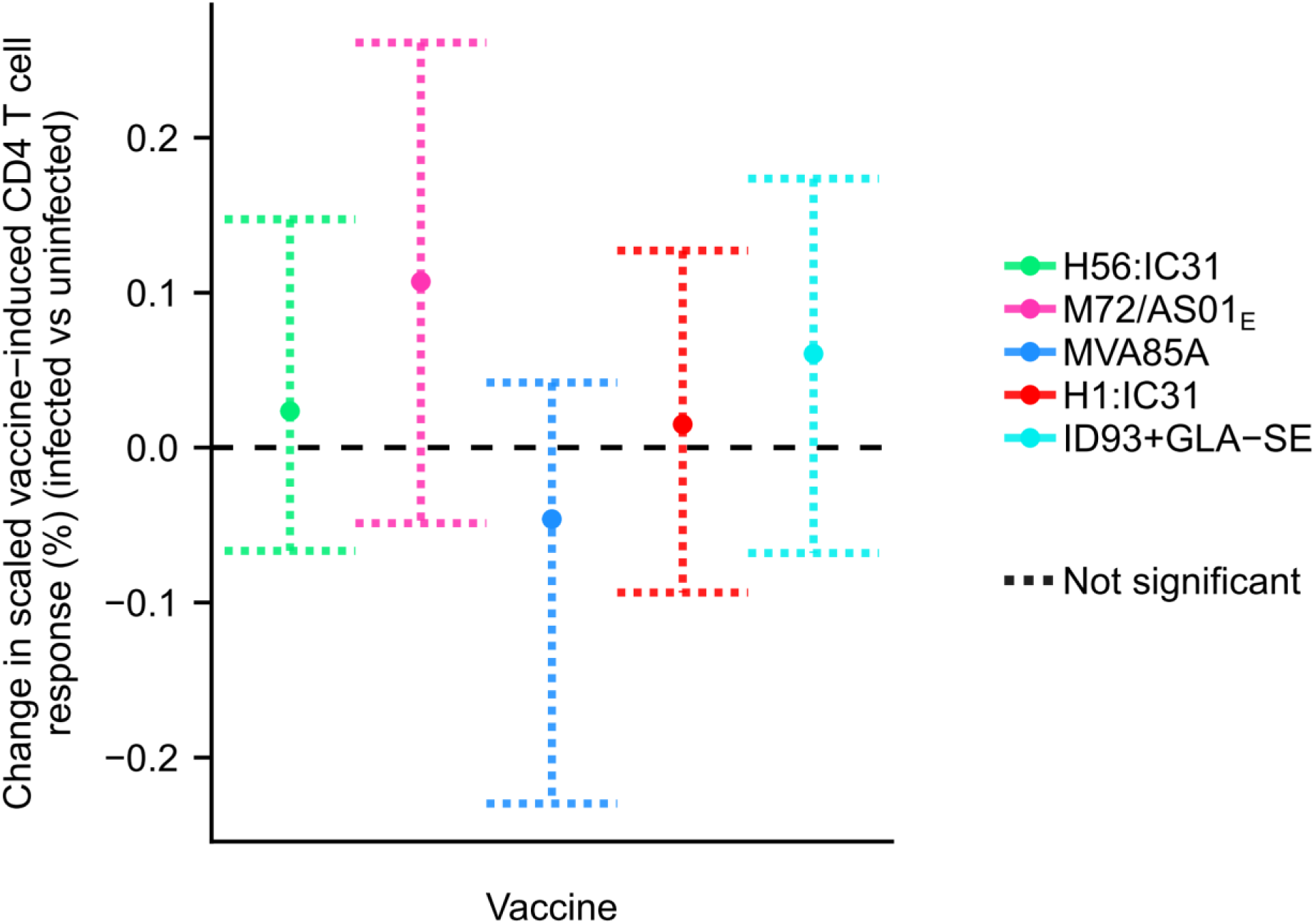
Effect of underlying *M.tb* infection status on vaccine-induced memory CD4 T cell response magnitudes. Differences between *M.tb*-infected and ‐uninfected individuals in vaccine-induced frequencies of antigen-specific memory CD4 T cell responses to antigens in each vaccine candidate (antigen-specific Th1 cytokine positive CD4 T cells at final trial time point minus pre-vaccination time point). Points denote sample trimmed means and error bars denote 95% CI. Solid error bar lines indicate responses that were significantly different between *M.tb*-infected and uninfected individuals given the same vaccine, after controlling the false discovery rate at 0.01. Dashed lines did not meet this significance criterion.

**S6 Fig.**
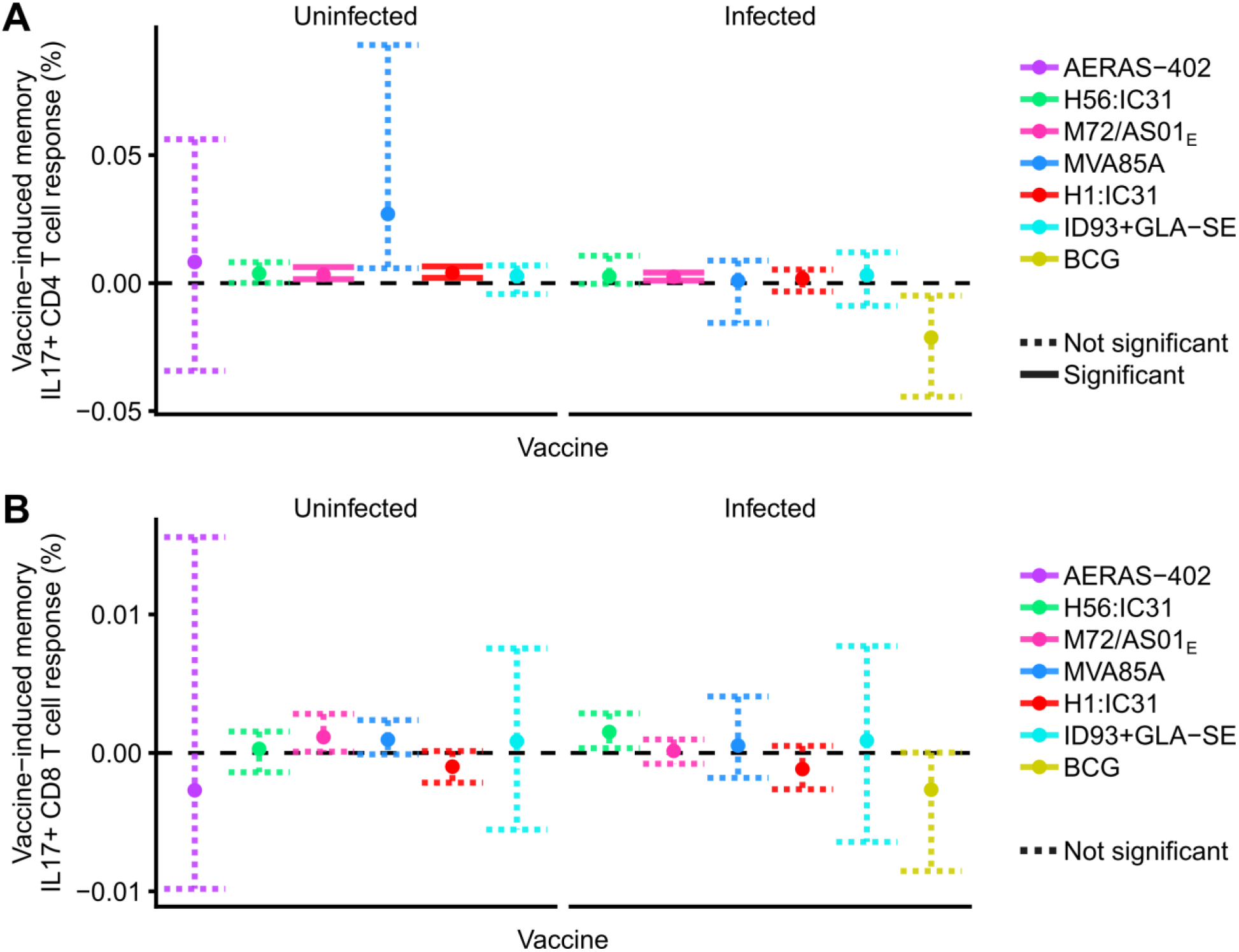
Vaccine-induced IL-17+ memory CD4 (A) and CD8 (B) T cell responses by vaccine and *M.tb* infection status. Frequencies of antigen-specific IL17-expressing CD4 or CD8 responses at the final time point in each trial, relative to the pre-vaccination frequencies (i.e. memory response minus pre-vaccination response). Points denote sample trimmed means, and error bars 95% CI. Solid error bar lines indicate responses that significantly exceeded 0% after controlling the false discovery rate at 0.01. Dashed lines did not meet this significance criterion.

**S7 Fig.**
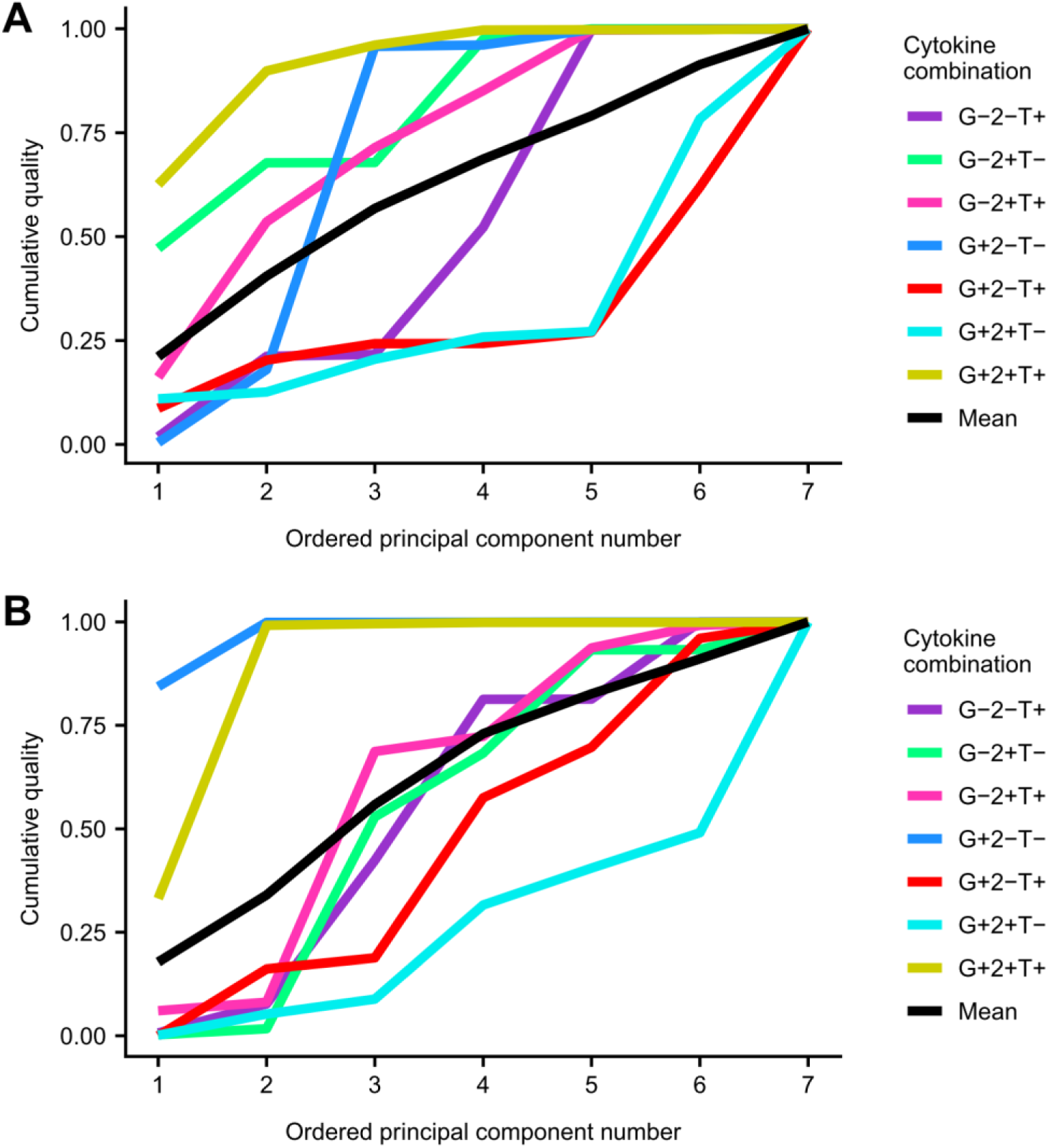
Cumulative axis predictivity of cytokine co-expression profiles for vaccine-induced memory CD4 T cells. Cumulative axis predictivity for k principal components represents the proportion of variation in the scaled vaccine-induced memory response for each cytokine combination captured by the first *k* principal components (see Materials and Methods for details).

**S8 Fig.**
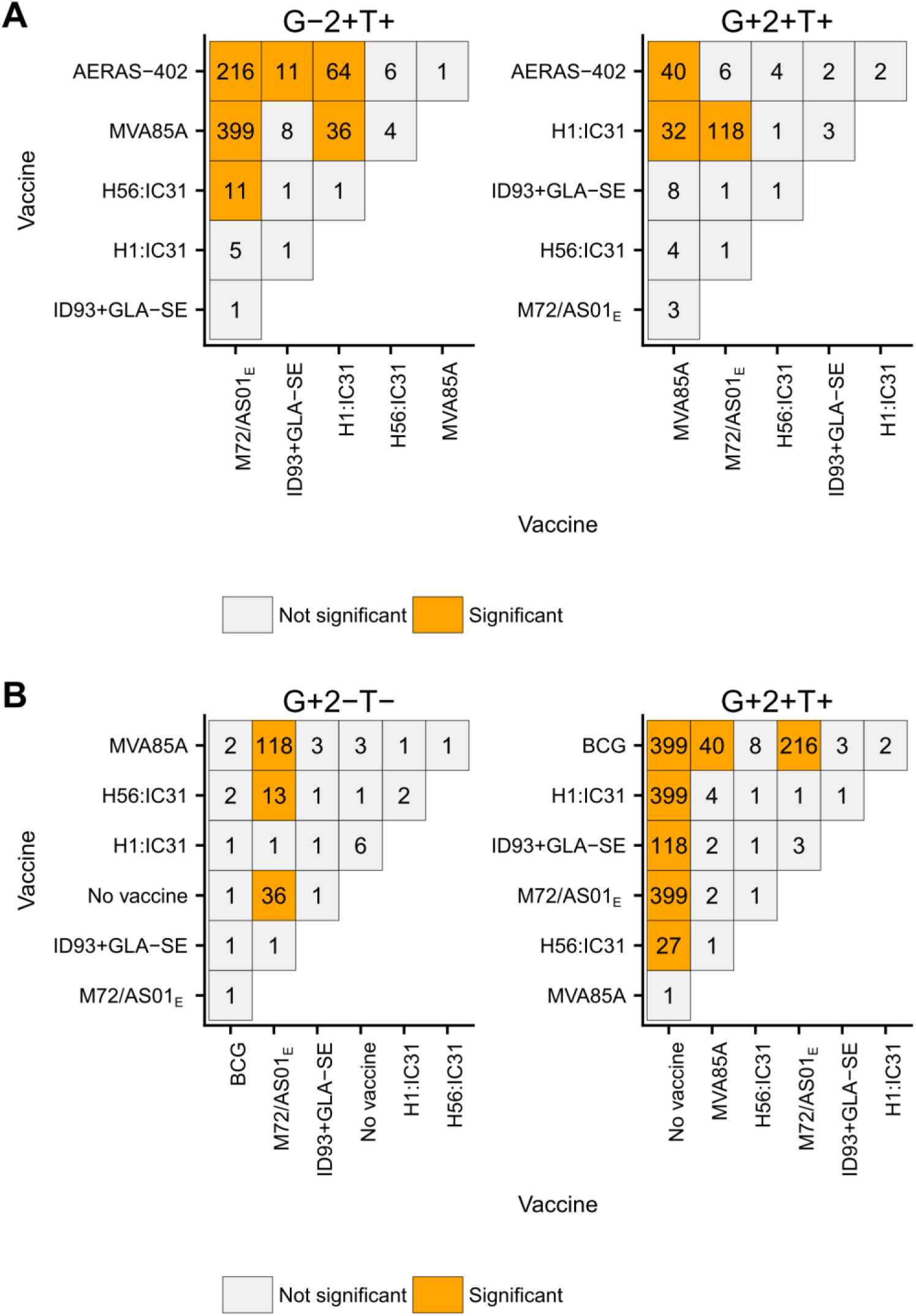
Pair-wise comparisons of cytokine co-expression profiles for vaccine-induced memory CD4 T cell responses between vaccines. Cells display maximum Bayes factors calculated based on the p-values from hypothesis tests for a difference between vaccines in scaled vaccine-induced memory CD4 T cell responses for certain cytokine combinations in *M.tb*-uninfected (A) and ‐infected (B) individuals (see Materials and Methods for details). The colour of the blocks indicates statistical significance of the two-sided bootstrap hypothesis test for a difference in population trimmed means, after controlling the false discovery rate at 0.05. An orange block means that the induced response for the vaccine below the block was significantly larger than the vaccine left of the block. A grey block indicates non-significance. For example, the figure shows that M72/AS01E produced statistically significantly higher TNF+IL-2+ CD4 T cell responses in *M.tb*-uninfected individuals than H56:IC31, but not ID93+GLA-SE.

**S9 Fig.**
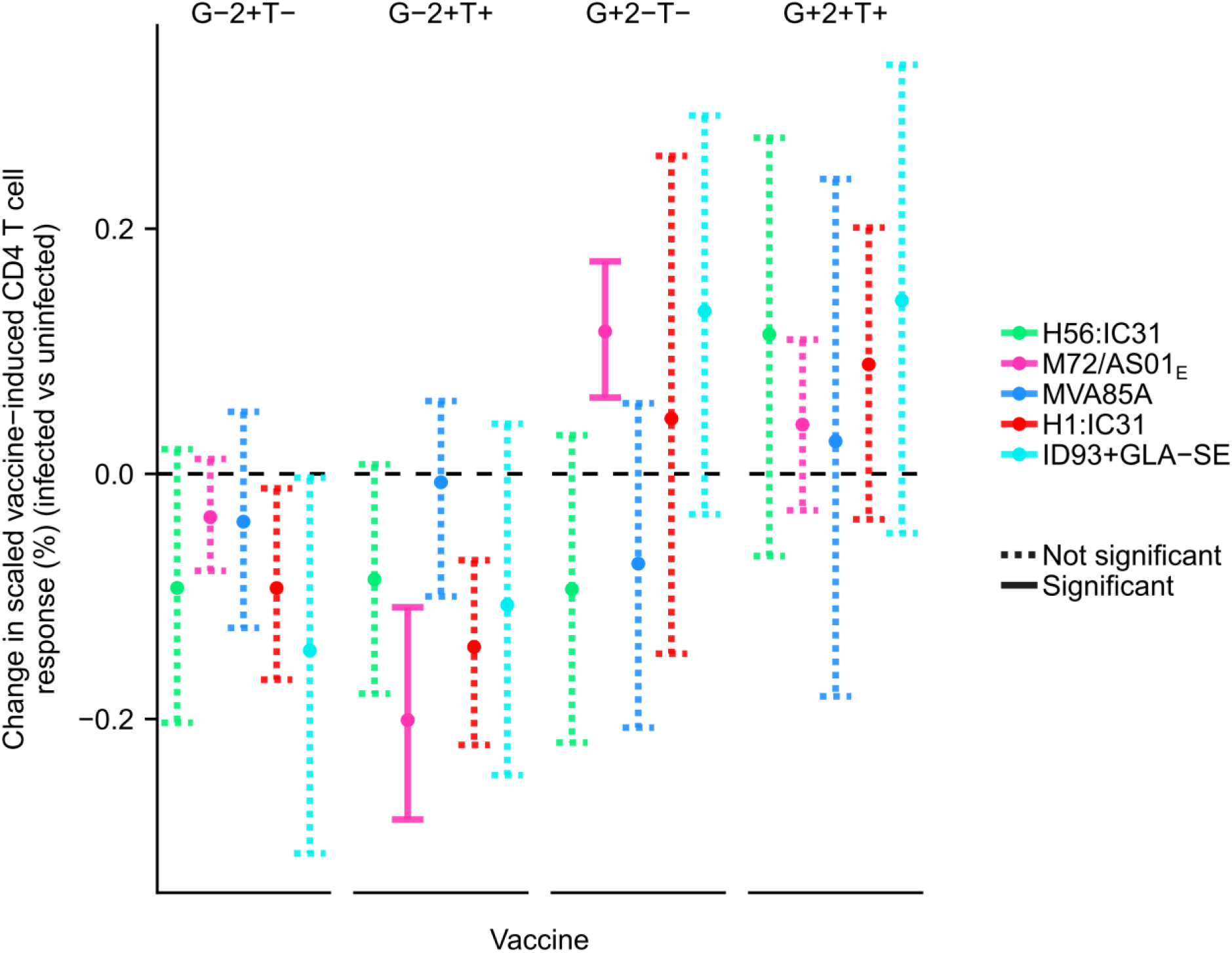
Effect of underlying *M.tb* infection status on vaccine-induced memory response profile for CD4 T cells for selected cytokine co-expression subsets. Differences between *M.tb*-infected and ‐uninfected individuals in scaled vaccine-induced frequencies of antigen-specific memory TNF+IL-2+, single IFNγ+, single IL-2+ and IFNγ+TNF+IL-2+ CD4 T cell responses to antigens in each vaccine candidate (see Materials and Methods for details). Points denote sample trimmed means and error bars denote 95% CI. Solid error bar lines indicate responses that were significantly different between *M.tb*infected and ‐uninfected individuals given the same vaccine, after controlling the false discovery rate at 0.01. Dashed lines did not meet this significance criterion.

**S1 Table.**
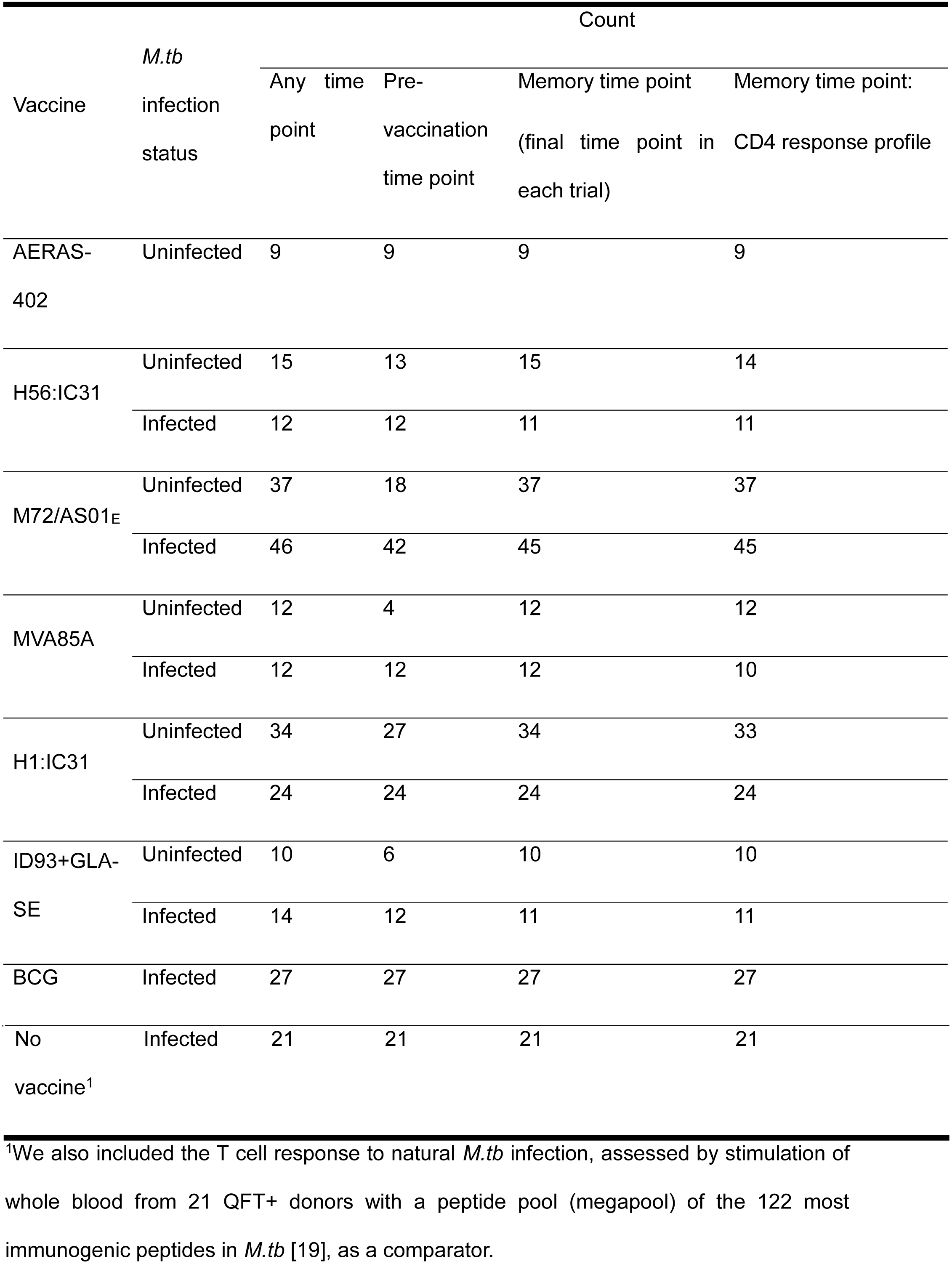
Sample sizes by time point for each vaccine trial.

